# Ubiquitin receptors are required for substrate-mediated activation of the proteasome’s unfolding ability

**DOI:** 10.1101/592378

**Authors:** Mary D. Cundiff, Christina M. Hurley, Jeremy D. Wong, Aarti Bashyal, Jake Rosenberg, Eden L. Reichard, Nicholas D. Nassif, Jennifer S. Brodbelt, Daniel A. Kraut

## Abstract

The ubiquitin-proteasome system (UPS) is responsible for the bulk of protein degradation in eukaryotic cells, but the factors that cause different substrates to be unfolded and degraded to different extents are still poorly understood. We previously showed that polyubiquitinated substrates were degraded with greater processivity (with a higher tendency to be unfolded and degraded than released) than ubiquitin-independent substrates. Thus, even though ubiquitin chains are removed before unfolding and degradation occur, they affect the unfolding of a protein domain. How do ubiquitin chains activate the proteasome’s unfolding ability? We investigated the roles of the three intrinsic proteasomal ubiquitin receptors - Rpn1, Rpn10 and Rpn13 - in this activation. We find that these receptors are required for substrate-mediated activation of the proteasome’s unfolding ability. Rpn13 plays the largest role, but there is also partial redundancy between receptors. The architecture of substrate ubiquitination determines which receptors are needed for maximal unfolding ability, and, in some cases, simultaneous engagement of ubiquitin by multiple receptors may be required. Our results suggest physical models for how ubiquitin receptors communicate with the proteasomal motor proteins.

---

Misfolded and damaged proteins, short-lived transcription factors and other regulatory proteins are all degraded by the Ubiquitin-Proteasome System (UPS) in eukaryotic cells ^1,2^. The typical signal for degradation is the attachment of a polyubiquitin chain to one or more lysine residues within the substrate by the action of E1, E2 and E3 enzymes. Ubiquitin is attached to substrate proteins through a covalent isopeptide bond between the C-terminus of ubiquitin and a lysine amino-group on the substrate. If a second ubiquitin is then attached to a lysine within the first ubiquitin, a polyubiquitin chain can begin to form. Depending on the E2 (or E3 for HECT- or RBR-type E3 ligases) involved, different lysine residues within ubiquitin can be used, leading to different polyubiquitin chain topologies ^3,4^.

The most common polyubiquitin chain linkages found in yeast are formed through K48 or K63 of ubiquitin ^5^. K48-linked chains are the canonical signal for proteasomal degradation. K63-linked chains, in contrast, are typically used in cellular trafficking and signaling, although for certain substrates and under certain conditions they can also function in proteasomal degradation ^6–8^. Both K48- and K63-linked chains function as degradation signals *in vitro*, although K63-linked chains are disassembled more rapidly by proteasomal deubiquitinases (DUBs) ^9,10^. Additional linkages and more complicated structures are also possible, and mixed-linkage and branched chains have been observed both *in vitro* and *in vivo*. Indeed, branched chains may enhance degradation of at least some substrates by increasing affinity of substrates for the proteasome or possibly allowing interaction with multiple ubiquitin receptors simultaneously ^11^.

Polyubiquitinated substrates interact with ubiquitin receptors on the 19S regulatory particle of the proteasome. The three known intrinsic proteasomal ubiquitin receptors are the ubiquitin interaction motif (UIM) of the Rpn10 subunit, the pleckstrin-like receptor for ubiquitin (pru) of Rpn13 (which is the entire subunit in yeast), and the T1 toroidal region of the Rpn1 subunit ^12–14^. These domains, like most ubiquitin-binding domains, are small and only interact with one or two ubiquitin molecules at a time ^15^. Mutation of a single ubiquitin receptor is well-tolerated by yeast, suggesting some level of functional redundancy, although some substrates may prefer one receptor to another ^15^. In fact, yeast with all three receptors mutated remain viable, and a proteasome that is mutant for all receptors is still capable of degrading some proteins, suggesting the potential for additional receptors that have yet to be discovered ^14^. It remains unclear why the proteasome contains such an array of ubiquitin-binding functionalities as well as what additional roles different ubiquitin receptors may play in the overall mechanism of protein degradation by the proteasome.

After binding of the polyubiquitin chain to the proteasome, an unstructured region of the substrate is engaged by the ATP-dependent motor proteins of the 19S regulatory particle (Rpt1-6 in yeast) ^16^. After engagement, the proteasomal ATPases begin pulling on the substrate, leading to first the removal of the polyubiquitin chain by the Rpn11 deubiquitinase ^17,18^, and subsequently to the unfolding of the substrate protein and its translocation into the 20S core particle, where it is hydrolyzed into short peptides ^19^ (Figure 1).

**Figure 1.**
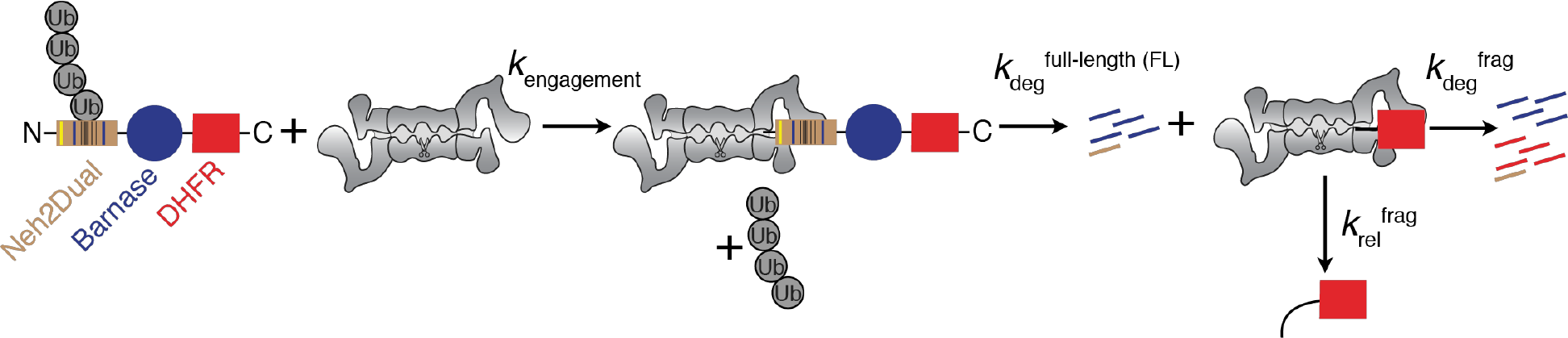
Schematic depiction of the assay. Substrate consists of an N-terminal Neh2Dual domain followed by a barnase domain and finally a C-terminal DHFR domain. Ubiquitinated substrate binds to the proteasome, is engaged by the proteasomal ATPases with concomitant removal of the polyubiquitin chain by Rpn11 (*k*_engagement_) and is degraded with the rate constant *k*_deg_^FL^, which represents unfolding the barnase domain and progressing to the point where DHFR is stalled at the entrance to the 19S regulatory particle. At this point the proteasome can either proceed to unfold and degrade DHFR (*k*_deg_^frag^) or release a stable DHFR fragment (*k*_rel_^frag^). The partitioning between the pathways is determined by the ratio of the rate constants for the two processes. For simplicity, non-productive deubiquitination of the substrate, which can occur in competition with engagement and initial degradation, is not shown.

Over the past several years, it has become increasingly clear that rather than just serving as the location of degradation, accepting all substrates with the appropriate polyubiquitin tag, the proteasome is intimately involved in the decision of whether or not to degrade a substrate ^20^. Deubiquitinases (DUBs) and ubiquitin ligases that associate with the proteasome compete to determine the residency time of a substrate before engagement, and the polyubiquitin chain itself can affect the peptidase and ATPase hydrolysis rates of the proteasome ^20,21^. We recently showed that the polyubiquitin modification on a substrate was surprisingly able to increase the ability of the proteasome to unfold and degrade a downstream domain of the substrate (increasing proteasomal processivity), even though the ubiquitin is presumably removed early in the degradation process, before the domain is unfolded (Figure 1) ^22^. Further, the details of substrate ubiquitination affected the proteasome’s unfolding ability. Substrates that were ubiquitinated by the Keap1/Cul3/Rbx1 E3 ligase complex (referred to as Keap1 for brevity), yielding mixed-linkage chains containing both K48- and K63-linkages, gave a higher unfolding ability than those that were ubiquitinated by Rsp5, which yielded exclusively K63-linked chains. Either mode of ubiquitination gave higher unfolding abilities than those seen with substrates targeted to the proteasome via ubiquitin-independent degrons. The mechanism by which ubiquitin conjugates activate the proteasome’s unfolding ability remains unknown, as does the mechanism for differential activation by Keap1- and Rsp5-ubiquitinated substrates. Here we show that proteasomal ubiquitin receptors are required for this substrate-mediated activation of the proteasome’s unfolding ability. These receptors can function redundantly, but more than one receptor is required for maximal activation with some substrates.

## Experimental procedures

### Constructs

Neh2Dual-Barnase-DHFR-His and the ubiquitin-independent substrate yO44-BarnaseL89G-DHFR-His were described previously; all barnase constructs were lysine-free ^22^. To destabilize barnase in the Neh2Dual construct, the L89G mutation ^23^ was introduced by oligo-directed mutagenesis to generate Neh2Dual-BarnaseL89G-DHFR-His. A 3C-Pro cleavage site between barnase and DHFR was introduced by oligo-directed mutagenesis. Individual lysines in DHFR were mutatated to arginine (as indicated) using oligo-directed mutagenesis in the Neh2Dual-BarnaseL89G-3CPro-DHFR-His construct, and a 35-amino acid lysine free linker derived from the pre-sequence to yeast cytochrome b_2_ (35ΔK) was introduced between the 3C-Pro site and DHFR (giving a 43-amino acid N-terminal unstructured region after cleavage) using Gibson assembly. Size standard constructs (Barnase-DHFR-His and DHFR-His) were derived from the above constructs using PCR methods. For bacterial overexpression, a Neh2Dual-Barnase-DHFR construct was moved into pE-SUMO, which provides an N-terminal His-SUMO tag, or into an MBP-containing expression vector, which provided an N-terminal MBP followed by an HRV 3C-Pro cleavage site. This second construct also had all cysteines mutated, and a single cysteine introduced into the linker between barnase and DHFR.

A plasmid expressing GST-Neh2Dual in bacteria under control of the T7 promoter was constructed by replacing vOTU from pOPINK-vOTU (Addgene plasmid # 61589, a gift from David Komander ^24^) using Gibson cloning.

A plasmid expressing ubiquitin with a cysteine inserted after the N-terminal methionine was created from ubiquitin-pET-3a ^25^ by oligo-directed mutagenesis.

Mutant ubiquitin receptors were expressed in yeast between the 5’ and 3’ UTRs of yeast Rpt1 in centromeric plasmids derived from Dp2 and Dp22, which were a gift from Dan Finley ^26^. Plasmids were constructed using traditional or Gibson cloning, and mutations were created using oligo-directed mutagenesis.

### Yeast strains

Yeast knockout strains were made in the background of strain YYS40 ^27^, which contains a 3XFLAG-tagged copy of Rpn11 to aid in proteasome purification. To make single gene knockout strains, the gene was replaced with a NatMX marker by PCR-directed homologous recombination ^28^. In the case of the essential gene Rpn1, a wild-type Rpn1 centromeric cover plasmid carrying a URA3 marker was used to cover the knockout, which was then replaced with a LEU2-containing plasmid with the Rpn1ΔT1 gene by plasmid shuffling using selection with 5-fluoroorotic acid (5-FOA). For non-essential genes, a centromeric plasmid containing the mutant ubiquitin receptor was transformed into the knockout strain to generate mutant proteasome. To create double ubiquitin receptor mutants, the single-mutant strain was switched from NatMX to KanMX resistance by PCR-directed homologous recombination, and the second gene was knocked out as above. To create a triple ubiquitin receptor mutant, a Cas9/gRNA construct targeting Rpn10 along with an Rpn10ΔUIM donor DNA were co-transformed into a ΔRpn13/ΔRpn1 strain containing an Rpn1ΔT1 LEU plasmid ^29^. After removal of the Cas9/gRNA plasmid by 5-FOA selection, an Rpn13-pru plasmid was transformed into the strain to create a strain containing Rpn1ΔT1, Rpn10ΔUIM and Rpn13-pru mutations. Strains are listed in **Supporting Table S1**.

### Proteasome purification

Yeast (*S. cerevisiae*) proteasome was purified from wild-type (YYS40) and mutant strains via a 3X-FLAG-tag on the Rpn11 subunit of the 19S regulatory particle essentially as described previously ^22^. Strains containing plasmids with mutant ubiquitin receptors were grown overnight in selective media before being transferred to YEPD media for larger scale growth. There was no detectable loss of plasmid during growth of the large-scale culture, as determined by comparing the number of viable colonies on YEPD and selective media at the time of harvesting. All proteasome preps showed primarily singly and doubly capped proteasome as determined by native gel analysis using the fluorogenic substrate Suc-LLVY-AMC ^30^ (**Supporting Figure S1**). Rpn10ΔUIM/Rpn13-pru proteasome was further purified by size exclusion chromatography in the presence of ATP on a Superose 6 column (Pharmacia). Pure fractions, as determined by native gel analysis, were pooled and concentrated using an Amicon Ultra 30 kDa cutoff centrifugal concentrator.

### Bacterial protein expression and purification

GST-Neh2Dual was overexpressed in BL21(DE3) cells in autoinducing media ^31^ at 30 °C and purified on glutathione agarose followed by gel filtration on a Sephacryl S200 column in 20 mM Tris, 150 mM NaCl, 1 mM DTT pH 7.5. His-SUMO-Neh2Dual-Barnase-DHFR was overexpressed in BL21(DE3) cells in autoinducing media ^31^ at 18 °C. Cells were resuspended in lysis buffer (50 mM sodium phosphate, 300 mM sodium chloride, 10 mM imidazole, pH 8.0), lysed by high-pressure homogenization, bound to a NiNTA column, washed, and eluted with increasing concentrations of imidazole. The protein was then dialyzed into 20 mM Tris, 150 mM NaCl, 1 mM DTT pH 7.5 in the presence of SUMO protease before being re-passed over the NiNTA column to remove the SUMO-tag and protease, and then was flash frozen and stored at −80 °C. MBP-Neh2Dual-Barnase-Cys-DHFR-His was expressed and purified identically, except that after elution from the NiNTA resin, protein was treated with DTT, ammonium sulfate precipitated, and labeled with sulfo-cyanine5 maleimide (Lumiprobe) according to published protocols ^32^. Free dye was removed by gel filtration on a Superdex 75 column (GE). Labeling was essentially quantitative as determined by absorbance at 280 and 646 nm. Cys-ubiquitin was purified via the same procedure as wild-type ubiquitin (except that 1 mM DTT was added to keep a reducing environment) ^22^. Purified cys-ubiquitin was then labeled with sulfo-cyanine3 maleimide (Lumiprobe) as described above. vOTU was purified as described previously ^22^.

### Substrate translation and ubiquitination

Radioactive protein substrates were *in vitro* translated and purified via their His tags on magnetic NiNTA beads (Cube-Biotech) as described previously ^22^. Neh2Dual substrates were then ubiquitinated and purified by spin size exclusion chromatography as described previously ^22^, while ubiquitin-independent substrates and size standards were used without additional purification. Substrates were flash-frozen and stored at −80 °C until use. To determine if ubiquitination occurred only on the degron, or on the DHFR domain, a substrate containing an HRV 3C protease site between the barnase and DHFR domains (Neh2Dual-BarnaseL89G-3Cpro-DHFR-6XHis) was ubiquitinated and purified before overnight cleavage with HRV 3C protease (4°C) followed by repurification on magnetic NiNTA beads with a denaturing wash (8 M Urea, 100 mM sodium phosphate, 100 mM TrisCl, 0.05% Tween-20, pH 8.0), a native wash (50 mM NaPhosphate, 300 mM NaCl, 40 mM imidazole, 0.05% Tween-20, 0.1 mg mL^−1^ bovine serum albumin (BSA), pH 7.4) followed by elution with elution buffer (50 mM sodium phosphate, 300 mM sodium chloride, 250 mM imidazole, 0.05% Tween-20, 0.1 mg mL^−1^ BSA, pH 8.0). Substrate was then incubated with 2 μM vOTU in UbiCREST buffer (final composition 62.5 mM Tris, 125 mM NaCl, 10 mM DTT, pH 7.5) ^33^. After incubation at 37 °C for 30 min, the reaction was quenched in SDS-PAGE loading buffer and analyzed by SDS-PAGE followed by phosphorimaging. The fraction modification on the DHFR domain was calculated by dividing the intensity of the untreated DHFR bands by the vOTU-treated bands.

GST-Neh2Dual was ubiquitinated as described previously ^22^ with modifications to account for the higher substrate concentrations. The Rsp5 ubiquitination reaction contained 166 nM E1, 5.88 μM UbcH7 (E2), 2.82 μM Rsp5 (E3), 4 mM ATP, 1 μM DTT, 4 μM substrate, and 1.33 mg mL^−1^ ubiquitin in Rsp5 ubiquitination buffer (25 mM TrisCl, 50 mM NaCl, 4 mM MgCl_2_ pH 7.5) and was incubated for 1.5 hours at 25 °C. The Keap1 ubiquitination reaction contained 130 nM E1, 2 μM UbcH5 (E2), 0.5 μM Cul3/Rbx1 (E3), 0.5 μM Keap1 C151S, 5 mM ATP, 4 μM substrate, and 1.45 mg mL^−1^ ubiquitin in Keap1 ubiquitination buffer (45 mM TrisCl, 100 mM NaCl, 10 mM MgCl_2_ pH 8.0) and was incubated for 1 hour at 25 °C. Ubiquitinated protein was then incubated with glutathione agarose beads in wash buffer (50 mM HEPES pH 6.8, 50 mM NaCl, 5% glycerol, 1 mM EDTA, 2 mM DTT, 0.05% Tween-20) for 1 hour at 4 °C, washed extensively with wash buffer, and then incubated with wash buffer containing 0.225 mg mL^−1^ HRV 3C protease for 30 minutes at room temperature to elute ubiquitinated Neh2Dual.

Neh2Dual-Barnase-DHFR was ubiquitinated identically to GST-Neh2Dual, but was then used for bottom-up mass spectrometry to identify ubiquitination sites without further purification. MBP-Neh2Dual-Barnase-Cys-DHFR-His was ubiquitinated identically, except that the reactions contained a mixture of 90% wild-type ubiquitin and 10% Cy3-labeled ubiquitin. The ubiquitination reactions were then incubated with amylose resin (NEB) for 1 hour at 4°C and washed with three times with 20 mM Tris-Cl, 200 mM NaCl, 1 mM EDTA, pH 7.5. The resin was then incubated with maltose elution buffer (20 mM Tris-Cl, 200 mM NaCl, 1 mM EDTA, 10 mM Maltose, pH 7.5) at room temperature for 1 hour to elute the ubiquitinated MBP-Neh2Dual-BarnaseΔK-His. The supernatant was then incubated with 0.15 mg mL^−1^ HRV 3C protease for 1 hour to remove the MBP domain.

### Middle-Down Mass Spectrometry

For each middle-down digest, 1 μg of polyubiquitinated substrate (GST-Neh2Dual ubiquitinated with either Keap1 or Rsp5) was incubated with 1 μg of trypsin for 4 hours at 37 °C in a solution of 50 mM ammonium bicarbonate having a total volume of 50 μL. LC-MS analysis was performed using a Dionex UltiMate 3000 nanoLC system (Thermo-Fisher) interfaced to a Thermo Orbitrap Lumos tribrid mass spectrometer (Thermo-Fisher) modified to perform ultraviolet photodissociation (UVPD) at 193 nm ^34^. 1 μL of each sample was injected without further processing. The chromatographic separation was performed in a trap-and-elute manner using analytical PicoTip (30 cm, 75 μm I.D.) and trapping IntegraFrit (3 cm, 100 μm I.D.) columns (New Objective) packed in-house with polymer reverse-phase resin (Agilent) with a 1000-Å pore size. The ubiquitin digest products were eluted using a linear gradient from 15% B to 55% B, where B was 99% acetonitrile, with 0.1% aqueous formic acid making up the remainder of the eluent.

Species eluting from the analytical column were introduced to the mass spectrometer by electrospray using a voltage of 2 kV. The mass spectrometer was operated in the top-speed mode with 7 seconds of MS2 data acquisition following each MS1 scan. Using a targeted instrument method, ions with a *m/z* ratio corresponding to the 12+ charge state of ubiquitin with 0 to 3 diglycine additions were isolated using the quadrupole then activated in the low-pressure ion trap by a single 1.8-mJ laser pulse. MS1 spectra were collected using a resolution of 60,000 with 3 microscans per recorded scan; MS2 spectra were collected with a resolution of 120,000 and 5 microscans per scan. The data were manually interpreted using Thermo Qual Browser and Prosight Lite software.

### Liquid Chromatography and Bottom-Up Mass Spectrometry

Ubiquitination sites were characterized using a bottom-up LCMS/MS strategy based on one reported by Gygi *et al*. in which glycine-glycine tagged peptides are produced upon trypsin proteolysis of ubiquitinated proteins ^35^. 5 μg of Neh2Dual-Barnase-DHFR substrate was incubated with 0.1 μg of trypsin for the control protein (not exposed to any ubiquitinating enzymes) or 0.4 μg of trypsin for the Rsp5-ubiquitinated protein in the presence of 50 mM ammonium bicarbonate at 37 °C for 16 hours. 2 μg of Keap1-ubiquitinated protein was reduced with 10 mM TCEP at 37 °C for 15 minutes before incubation with 0.5 μg of trypsin in presence of 50 mM ammonium bicarbonate at 37 °C for 16 hours. Trypsin was deactivated by addition of 0.5% formic acid, and the samples were cleaned up using Pierce C18 spin columns (Thermo Fisher Scientific) according to the procedure described by the manufacturer.

Each digest was separated using an Ultimate 3000 RSLCnano liquid chromatography system fitted with a New Objective IntegraFrit trap column (3.5 cm, 100 μm I.D.) and PicoFrit analytical column (20 cm, 75 μm I.D.), and the eluent was analyzed with a Thermo Orbitrap Lumos tribrid mass spectrometer. The columns were packed in-house using UChrom C-18 stationary phase (120 Å, 3 μm particles for the trap column and 120 Å, 1.8 μm particles for the analytical column). Tryptic digests were constituted in 0.1% formic acid, and approximately 250 ng of each digest was preconcentrated on the trap column at 5 μL/min for 5 min with 2% acetonitrile and 0.1% formic acid. The mixtures were then separated on an analytical column with a linear gradient from 2% B to 40% B over a period of 50 min at a flow rate of 300 nL/min. Mobile phases for separation on the analytical column consisted of 0.1% formic acid in water as solvent A and 0.1% formic acid in acetonitrile as solvent B.

For nanospray, 2 kV was applied at a precolumn liquid voltage junction. Automated gain control targets were 500000 for MS1 and 50000 for MS2, and the maximum ion time was 100 ms for MS1 and 150 ms for MS2. 2 microscans were collected for both MS1 (*m/z* 400-2000, 60K resolution) and MS2 (*m/z* 220-2000, 30K resolution) spectra. Peptide ions were activated by HCD using a normalized collision energy (NCE) of 30%. Data acquisition was performed at top-speed mode using a cycle time of 5 s. The dynamic exclusion was set to exclude a precursor *m/z* value after selection 3 times for the exclusion duration of 45 s.

### Database Searching

LC-MS/MS data was analyzed using Byonic™ by Protein Metrics Inc. (v3.1.0). All spectra were searched against a FASTA file containing the sequences of ubiquitin, trypsin, pDAK443, and Keap1. All searches employed a 10-ppm mass tolerance for both precursor and fragment ions. Fully specific searches were performed with up to 3 missed cleavages with 1% false discovery rate. Diglycine (+114.0429 Da) and LRGG (+383.2281 Da) tags were included in each search as a common modification at lysines. A PEP2D score of 0.001 was used as a cutoff margin for assigning the modified peptides. All spectra are archived and available at: https://repository.jpostdb.org/ and accession numbers are PXD010803 for ProteomeXchange and JPST000481 for jPOST.

### Degradation assay

Degradation assays were conducted with proteasome in great excess of substrate (100 nM proteasome and trace radiolabeled substrate), essentially as described previously ^22^, although time courses ranged from two to nine hours depending on the observed rate constant for degradation.

### Curve fitting, modeling, and data analysis

Dried gels were phosphorimaged using a Typhoon FLA9500 (GE). Gels were quantified and data was analyzed as previously described ^22,36^ using ImageJ (NIH). Time courses for the full-length protein and the DHFR-containing fragment from at least four assays were globally fit to single exponentials in Igor Pro (Wavemetrics) to determine the observed rate of degradation (*k*_obs_) and the amplitudes of degradation and fragment formation. These amplitudes were then used to calculate the unfolding ability (U) using Equation 1 (see Results). Statistical comparisons were made using a two-tailed Student’s T-Test.

### Peptidase assay

Proteasome (10 nM) was mixed with 1 mM ATP, an ATP-regeneration system consisting of 25 mM creatine phosphate and 0.1 mg mL^−1^ creatine kinase, and 50 μM substrate (Suc-LLVY-AMC, Z-LLE-AMC or Ac-RLR-AMC) in a buffer consisting of 50 mM Tris•Cl pH 7.5, 5 mM MgCl_2_, 2.5% glycerol, 2 mM DTT and 1% final DMSO. Proteasome inhibitors (100 μM each bortezomib, MG-132 and either 1,10-phenanthroline or epoxomicin) were optionally added. The appearance of AMC was monitored using excitation at 353 nm and emission at 442 nm on a ClarioSTAR Omega platereader and the linear portion of the curve was fit to a line to determine rates of reaction. Reactions with substrate (and inhibitors, as needed) but lacking proteasome were used to determine background rates.

## Results

### Proteasome lacking Rpn10 has difficulty unfolding substrate proteins

The ubiquitin receptor Rpn10 has previously been shown to be essential for degradation of a difficult-to-unfold GFP substrate by reconstituted proteasome ^37^. We therefore sought to determine if Rpn10 contributed to the ubiquitinated substrate-dependent activation of the proteasome’s unfolding ability. The model substrate we use to measure unfolding ability ^22^ contains an N-terminal Neh2Dual degron, which can be ubiquitinated by either Keap1 or Rsp5, followed by an easy-to-unfold lysine-free barnase domain and a C-terminal dihydrofolate reductase (DHFR) domain (Figure 1). After engaging the Neh2 domain and removing the polyubiquitin modification (*k*_engagement_), the proteasome proceeds along the substrate, first unfolding and degrading the barnase domain (*k*_deg_^FL^) and then encountering the DHFR domain (typically stabilized by the addition of NADPH). At this point, the proteasome either unfolds and degrades the resulting DHFR-containing fragment or, if the substrate slips out of the degradation channel, irreversibly releases it. The ratio of the degradation to release rates (the unfolding ability U) can be measured most easily by comparing the amount of full-length protein that is fully degraded by the proteasome with the amount of DHFR-containing protein fragment that is created by incomplete degradation of the full-length protein (Figure 1; **Equation 1**) ^22,36^.

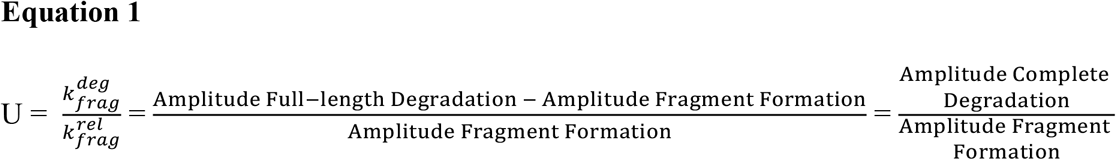

Control experiments in which the order of the DHFR and barnase domains are reversed confirm that degradation proceeds from the N-terminus of the protein, as DHFR stabilized by methotrexate (MTX) is able to protect barnase from degradation (**Supporting Figure S2**) ^38–40^.

One advantage of this tool is that the unfolding ability we measure is independent of the concentration of proteasome or substrate, as it is a measurement of the partitioning of the proteasome-DHFR fragment between two possible fates, degradation and irreversible release ^36^. Thus, when comparing different batches of proteasome or different mutants, errors in concentration determination will not affect the measured unfolding abilities. Similarly, differences in the deubiquitination of Keap1-versus Rsp5-ubiquitinated substrates, should they occur, may affect the total extent of degradation of full-length protein, but will not affect the ratio of protein that is partially degraded versus fully degraded.

When we conducted unfolding ability assays using proteasome purified from yeast lacking the Rpn10 protein, we found that degradation was qualitatively different compared to that observed with wild-type proteasome. Keap1-ubiquitinated substrate could hardly be degraded by ΔRpn10 proteasome (Figure 2A). Rsp5-ubiquitinated substrate was degraded, albeit much less efficiently than by wild-type proteasome (Figure 2B; wild-type degraded 85% of the substrate in 2 hours while ΔRpn10 degraded only 50%). Additionally, we found that the mutant proteasome produced higher molecular weight fragments of the substrate than wild-type, suggesting that the proteasome sometimes stalls after partially degrading Neh2 but before unfolding barnase (Figure 2B). As similar amounts of fragment were formed even though less of the full-length substrate was degraded, formation of this Barnase-DHFR-containing fragment and of a smaller DHFR-containing fragment occurred with a greater frequency with ΔRpn10 proteasome than with wild-type proteasome, and complete degradation was less frequent (Figure 2C). Thus, it appears that ΔRpn10 proteasome has more difficulty unfolding substrates than wild-type proteasome.

**Figure 2.**
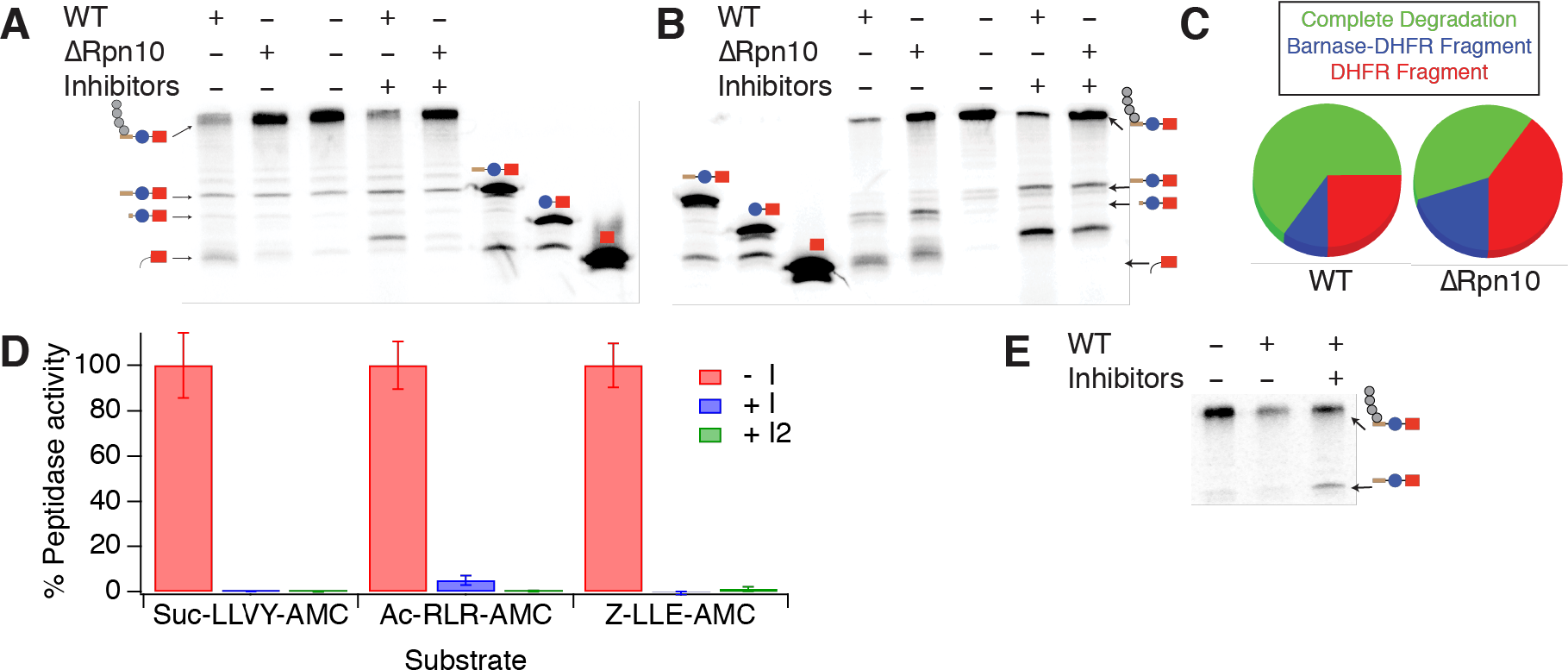
ΔRpn10 proteasome has a processivity defect. **A**, **B**) Degradation of trace radiolabeled Keap1-(**A**) or Rsp5-ubiquitinated substrate (**B**) by 100 nM wild-type or ΔRpn10 proteasome (2 hours). Fragments containing either DHFR plus a small tail or Barnase-DHFR plus a small tail were identified relative to size standard control constructs (Neh2Dual-Barnase-DHFR, Barnase-DHFR and DHFR). Proteasome inhibitors (100 μM each MG-132, Bortezomib and 1,10-phenanthroline) were added as indicated. Based on quantification, less than 10% of the full-length Keap1-ubiquitinated protein was degraded by ΔRpn10 proteasome. Note that similar or greater levels of fragment were formed with ΔRpn10 and the Rsp5-ubiquitinated substrate despite a far lesser extent of degradation. **C**) Quantification (based on two replicate experiments) of the relative frequency of complete versus partial degradation for wild-type versus ΔRpn10 proteasome with the Rsp5-ubiquitinated substrate. Protein that did not undergo degradation was ignored. **D**) Peptidase activity in the absence (red) or presence of proteasome inhibitors as in B (blue; I) or with 100 μM epoxomicin replacing 1,10-phenanthroline (green; I2). The first cocktail reduces Suc-LLVY-AMC (chymotrypsin-like) activity to 0.1% (p = 0.07 when comparing proteasome plus inhibitors to background reaction), reduces Ac-RLR-AMC (trypsin-like) activity to 5% of uninhibited rates (p = 0.001), and eliminates Z-LLE-AMC (peptidyl glutamyl hydrolase-like) activity (at or below background rates). With the second cocktail, all rates are indistinguishable from background. Error bars are the SEM of 4 replicate assays. **E**) Degradation of trace radiolabeled Rsp5-ubiquitinated substrate by 100 nM wild-type proteasome in the absence or presence of 100 μM each MG-132, Bortezomib, and epoxomicin. No NADPH was added to stabilize DHFR in this experiment.

To confirm that degradation was due to the proteasome, we tested the effect of proteasome inhibitors on the reaction (Figure 2B). The addition of proteasome inhibitors (100 μM MG-132, bortezomib and 1,10-phenanthroline) reduced the extent of degradation (from 85% to 70% for wild-type with the Rsp5 substrate and from 50% to 30% for ΔRpn10). Although we were initially puzzled that degradation was not completely inhibited, peptidase assays showed that the proteasome retained 5.1 ± 0.8% of its trypsin-like activity (Figure 2D, blue). Given the single-turnover-like conditions used (proteasome in excess over trace radiolabeled substrate) only a small fraction of the proteasome needs to be active for engagement to occur and degradation to be initiated, and it has been shown previously that proteasomes with multiple active sites inhibited retain the ability to degrade some substrates ^41^. The fragments resulting from partial degradation in the presence of inhibitors were larger than in their absence, suggesting that potential substrate cleavage sites closer to the DHFR or barnase domain are no longer cleavable. When we replaced 1,10-phenanthroline with epoxomicin in our proteasome inhibitor cocktail, the trypsin-like activity was reduced to 0.3 ± 0.3% (Figure 2D, green), and the extent of degradation was reduced to ~25% for wild-type proteasome (Figure 2E).

Although the inability of ΔRpn10 proteasome to unfold barnase suggests a major processivity defect, it also makes determination of an unfolding ability problematic. We therefore destabilized barnase by mutagenesis, which prevented the formation of the barnase/DHFR-containing fragment but not the DHFR-only fragment (Figure 3A, B) ^22^. Quantification of these degradation assays allowed us to determine the extent of degradation of the full-length substrate and the extent of formation of the DHFR-containing protein fragment, and thus the unfolding ability (U) (Figure 3C). U was approximately 2-fold lower for ΔRpn10 proteasome than for wild-type with either the Keap1 or Rsp5-ubiquitinated substrate, suggesting that Rpn10 is involved in ubiquitin-dependent activation of the proteasome’s unfolding ability. Unsurprisingly, there was little difference in unfolding ability (or rate of degradation) when using a ubiquitin-independent substrate where the Neh2Dual degron was replaced with the N-terminus of yeast ornithine decarboxylase (Figure 3); this substrate had previously been shown to be degraded with a very low unfolding ability ^22^. ΔRpn10 proteasome still unfolded and degraded weakly folded substrates, as the omission of NADPH led to highly processive degradation of ubiquitinated substrates, with no DHFR fragment formed (**Supporting Figure S3**). However, as Rpn10 contacts elements of both the proteasome base and lid subcomplexes and is important for their stability ^42–44^, it remained unclear if ubiquitin binding to Rpn10 was required for activation of the proteasome’s unfolding ability, or if instead Rpn10 was playing a structural or conformational role independent of its ability to bind ubiquitin.

**Figure 3.**
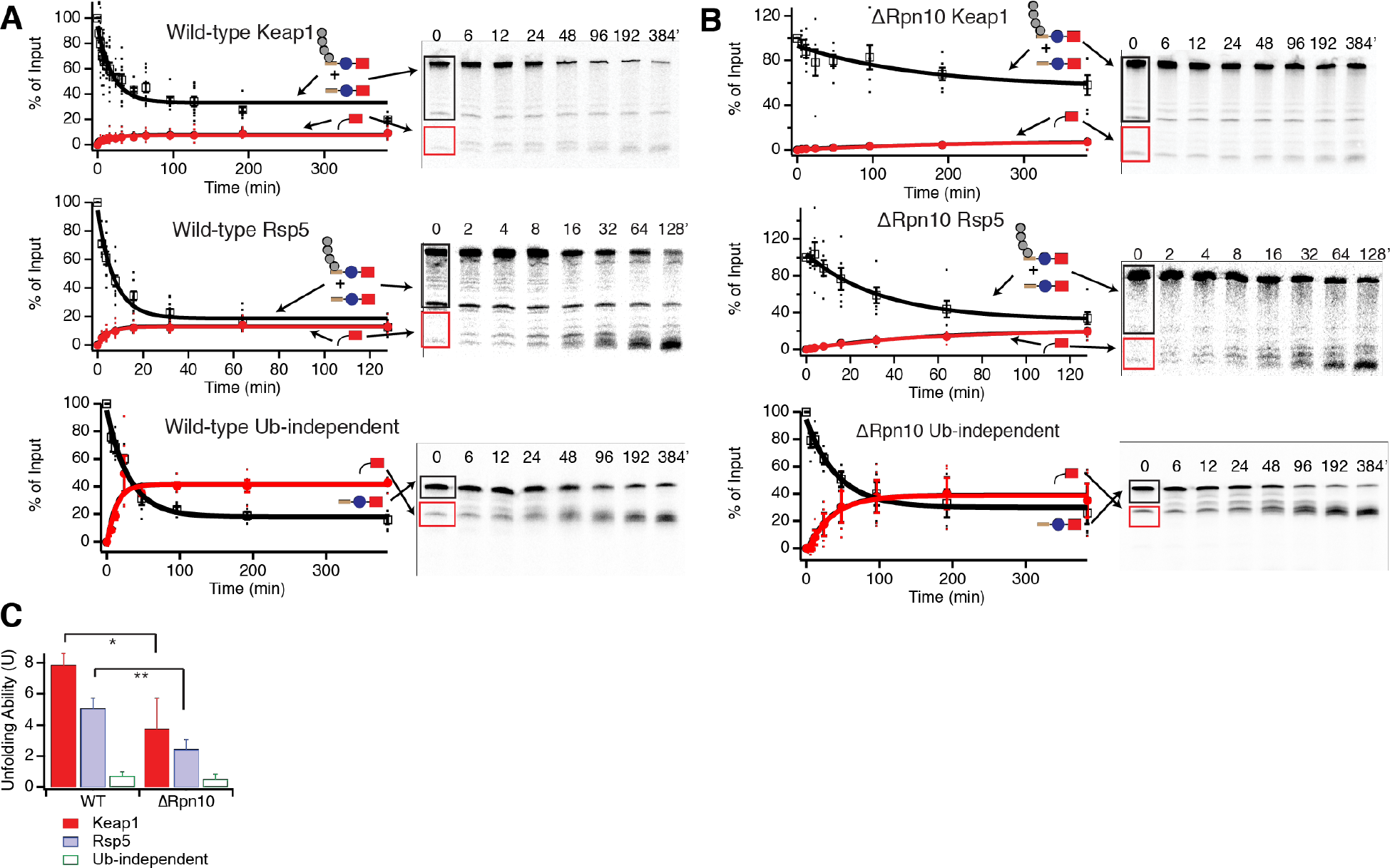
Unfolding abilities of wild-type versus ΔRpn10 proteasomes. **A**, **B**) Degradation of trace radiolabeled Keap1- or Rsp5-ubiquitinated substrate or ubiquitin-independent substrate (yO44-Barnase-DHFR-His) with a destabilized barnase domain by 100 nM wild-type (**A**) or ΔRpn10 (**B**) proteasome. Black box shows region of the gel containing full-length protein with or without ubiquitination. Red box shows the region of the gel containing the DHFR fragment. The amounts of full-length protein (open squares) and DHFR fragment (red circles) are shown as a percentage of the ubiquitinated full-length substrate presented to the proteasome at the beginning of the reaction (or the full-length protein for the ubiquitin-independent substrate); the full-length protein is quantified as the sum of ubiquitinated and non-ubiquitinated full-length species so any deubiquitination is not misinterpreted as degradation. Dots are results from individual experiments, and error bars represent the SEM of 6-22 experiments. Curves are global fits to single exponentials. Sample gels are shown on the right. **C**) Unfolding abilities (U) calculated from the curve fits shown in **A** and **B**. Error bars are the SEM propagated from curve fitting the collected data sets. * indicates p ≤ 0.05, ** indicates p ≤ 0.01 relative to wild-type.

### Rpn13 ubiquitin binding is required for maximal proteasomal processivity

To more incisively probe the contributions of individual ubiquitin receptors to proteasomal processivity without introducing gross structural or other functional changes to the proteasome, we made previously characterized point mutations that prevent ubiquitin binding to Rpn1 (Rpn1ΔT1: D541A/D548R/E552R) ^14^, Rpn10 (Rpn10ΔUIM: L228A/M230A/L232A) ^12^, or Rpn13 (Rpn13-pru: E41K/E42K/L43A/F45A/S93D) ^14^. Mutant proteasome containing only two functional ubiquitin receptors was then purified and assayed using the three substrates described above (Figure 4A, **Supporting Figure S4, Supporting Table S2**).

**Figure 4.**
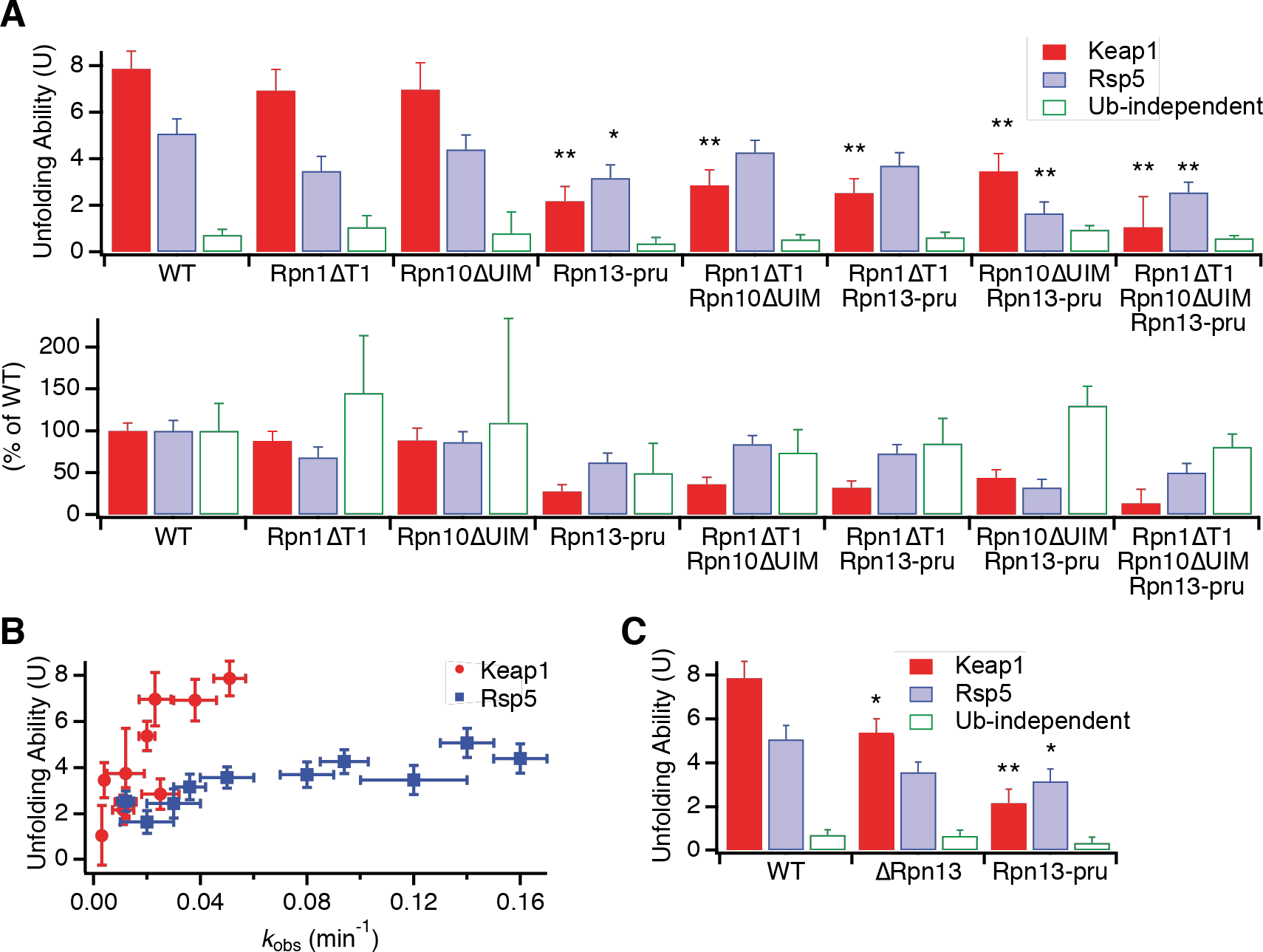
Ubiquitin receptor mutants have varied effects on proteasomal unfolding ability. **A**) Top: Effect of point mutants in ubiquitin binding domains on proteasomal unfolding ability, as measured using Keap1 (red) or Rsp5 (blue) ubiquitinated substrates or a ubiquitin-independent substrate (green). Error bars are the SEM propagated from curve fitting 4-22 collected data sets. * indicates p ≤ 0.05, ** indicates p ≤ 0.01 relative to wild-type. Bottom: unfolding abilities of mutants as normalized to wild-type proteasome. **B**) Unfolding ability (From panel A above) versus observed rate constant for disappearance of full-length protein (*k*_obs_) for Keap1 (red circles) and Rsp5 (blue squares) ubiquitinated substrates (from curve fits in Figure 3A and **Figure S4**). **C**) Rpn13 deletion (ΔRpn13) has a smaller effect on proteasomal unfolding ability than Rpn13 point mutations that prevent ubiquitin binding (Rpn13-pru). Error bars are the SEM propagated from curve fitting 4-22 collected data sets. * indicates p ≤ 0.05, ** indicates p ≤ 0.01 relative to wild-type.

For the Keap1 substrate, the Rpn1ΔT1 mutation had little effect and the Rpn10ΔUIM mutation reduced the observed rate constant for degradation (*k*_obs_) by ~2-fold but had no significant effect on unfolding ability. However, the Rpn13-pru mutation both substantially slowed degradation (~5-fold reduction in *k*_obs_) and reduced the unfolding ability from 7.9 ± 0.8 to 2.2 ± 0.6, closer to the level seen with a ubiquitin-independent substrate incapable of activating the proteasome’s unfolding ability (U = 0.7 ± 0.2).

For the Rsp5 substrate, which is less effective at activating the proteasome’s unfolding ability, the effects of all mutations were less substantial, with only the Rpn13-pru mutation giving a statistically significant (but smaller) reduction in unfolding ability (U = 5.1 ± 0.6 for wild-type, 2.7 ± 0.3 for Rpn13-pru) or in the observed rate constant for degradation (~3-fold reduction. As expected, none of the mutations had any statistically meaningful effect on the ability of the proteasome to unfold a ubiquitin-independent substrate, although it is difficult to accurately quantify unfolding abilities below 1, which corresponds to half of full-length protein that is degraded being converted into fragment, as small changes in the amount of fragment formed will translate into relatively large changes in the unfolding ability.

Thus, it appears that ubiquitin binding to Rpn13, but not Rpn10 or Rpn1, is necessary for maximal proteasomal activation by ubiquitinated substrates. The large effect of removing the entire Rpn10 subunit versus the minimal effect of preventing Rpn10 ubiquitin binding suggests a conformational or structural role for Rpn10.

### Proteasome with only one functional ubiquitin receptor reveals cooperation between receptors and functional redundancy

Mutating individual receptors showed that ubiquitin binding by Rpn13 was necessary for full proteasomal activation by Keap1-ubiquitinated substrates and was also important for unfolding of Rsp5-ubiquitinated substrates. To determine if Rpn13 was sufficient, we made double-mutants containing only one functional ubiquitin receptor (Figure 4A, **Supporting Figure S4, Supporting Table S2**). As expected, none of these mutations significantly affected the processivity or rate with which the proteasome unfolded a ubiquitin-independent substrate.

For the Keap1-ubiquitinated substrate, all of the double-mutants had low unfolding abilities of U ~ 2-4. Intriguingly, it appears that Rpn13 ubiquitin binding is not sufficient for a high unfolding ability, as the Rpn1ΔT1/Rpn10ΔUIM double-mutant, which still contains functional Rpn13, had an unfolding ability similar to the other double-mutants or the Rpn13-pru single-mutant. Therefore, two ubiquitin receptors are required for maximal proteasomal processivity with the Keap1-ubiquitinated substrate.

For the Rsp5-ubiquitinated substrate, both the Rpn1ΔT1/Rpn10ΔUIM and Rpn1ΔT1/Rpn13-pru double-mutants had unfolding abilities only slightly reduced from wild-type^*^, while the Rpn10ΔUIM/Rpn13-pru double-mutant had a greatly reduced unfolding ability (U = 1.7 ± 0.5). Thus, it appears that either Rpn10’s or Rpn13’s ability to bind ubiquitin is sufficient for some or most of the proteasomal activation achieved by Rsp5-ubiquitinated substrates, but proteasome containing only functional Rpn1 is not able to be activated. The relatively high unfolding ability of proteasome containing just a single ubiquitin receptor with the Rsp5-ubiquitinated substrate stands in contrast to results with the Keap1-ubiquitinated substrate, where two receptors were required for maximal activation. Thus the Keap1-ubiquitinated substrate has a higher overall unfolding ability but also needs to engage with two different receptors simultaneously while the Rsp5-ubiquitinated substrate is able to engage with either Rpn13 or Rpn10, albeit with some preference for Rpn13.

When a triple-mutant deficient in all known ubiquitin receptors was assayed, unfolding abilities with both the Keap1 and Rsp5-ubiquitinated substrate remained very low, and there was no change in U for the ubiquitin-independent substrate. Although degradation of ubiquitinated substrates was slowed substantially (~10-fold for either Keap1 or Rsp5-ubiquitinated substrates, versus no effect on the ubiquitin-independent substrate), it was not completely abrogated, suggesting the presence of at least one additional ubiquitin receptor, which might play a small role in activating the proteasome’s unfolding ability.

An alternative explanation for the decreased unfolding abilities we observe would be low levels of free 20S proteasome that are capable of slowly degrading unstructured regions within the substrate, but are unable to degrade the DHFR domain ^45^. If contaminating 20S contributed to degradation and DHFR fragment formation for mutants with lower overall degradation rates, we would be unable to accurately determine U for the 26S proteasome. However, several pieces of evidence suggest 20S contamination is not an issue. First, although our preps did contain some free 20S proteasome (**Supporting Figure S1**), the amount of 20S did not correlate with the observed unfolding ability. For example, the Rpn10ΔUIM/Rpn13-pru double-mutant had far more free 20S proteasome (~7%) than the triple-mutant (<1%), but they had very similar observed rates of reaction and unfolding abilities. Second, although there was a correlation between the observed rate constant for degradation and U (Figure 4B), it was not absolute – for example, both Rpn10ΔUIM (U = 7 ± 1 for Keap1) and Rpn1ΔT1/Rpn10ΔUIM (U = 2.9 ± 0.7 for Keap1) were degraded with rate constants of ~0.025 min^−1^, only a two-fold reduction from wild-type. Further, the overall dependence of U on the observed rate constant was much steeper for Keap1 than Rsp5, with different mutants having different effects on Keap1- and Rsp5-ubiquitinated substrates. Indeed, the Rpn10ΔUIM and Rpn1ΔT1/Rpn10ΔUIM mutants had indistinguishable unfolding abilities with Rsp5-ubiquitinated substrates (U = 4.4 ± 0.6 and 4.3 ± 0.5). Third, 20S proteasome (or other contaminating proteases) should not be able to degrade well-folded domains such as DHFR. In the absence of NADPH, which stabilizes DHFR but is not essential for its folding, both wild-type and mutant proteasomes were able to degrade ubiquitinated substrates without (for Keap1-ubiquitinated substrates) or with very low levels (for Rsp5-ubiquitinated substrates) of DHFR-containing fragment being produced, suggesting that mutant proteasomes retain the ability to unfold and degrade more weakly folded domains (**Supporting Figure S3**). Indeed, we had previously shown that although the proteasome was hardly able to unfold and degrade a ubiquitin-independent substrate containing DHFR stabilized by NADPH, it was able to easily degrade DHFR in the absence of NADPH ^22^. Fourth, to exclude the possibility of other contaminating proteases affecting our results, we repurified Rpn10ΔUIM/Rpn13-pru proteasome via gel filtration. The resulting proteasome gave equivalent unfolding abilities (**Supporting Table S2**), although the level of free 20S was unaffected, suggesting the mutations may have some direct effects on the assembly state of the proteasome.

Ubiquitinated substrates had previously been shown to activate the ATPase activity of the proteasome by binding to Ubp6 ^46^, a proteasome-associated deubiquitinase that can either accelerate or prevent degradation, depending on the circumstances ^47–49^. However, rates of ATP hydrolysis are not necessarily directly connected to rates of protein unfolding and degradation, and might not be maintained after engagement and removal of the ubiquitin chain. Indeed, our substrates had little to no ability to enhance the ATPase activity of the proteasome (data not shown), suggesting that either ubiquitin receptors work to increase unfolding ability independent of Ubp6 or that, if Ubp6 is involved, the previously observed Ubp6-linked increase in ATP hydrolysis rates is not required for enhancing the ability to unfold proteins.

### Rpn13 may repress the proteasome until ubiquitin binds

In yeast, Rpn13 is a single pleckstrin-homology-domain protein whose only known function is ubiquitin and ubiquitin-like protein binding. Rpn13 is at the apical tip of the 19S regulatory subunit ^43^, so we therefore reasoned that removing the entire Rpn13 subunit should give similar results to merely removing its ability to bind ubiquitin. However, in contrast to the large rate and unfolding ability defects seen with Rpn13-pru, ΔRpn13 proteasome had only a modest reduction in unfolding ability with the Keap1-ubiquitinated substrate (U = 5.4 ± 0.6 for ΔRpn13 versus 7.9 ± 0.8 for wild-type and 2.2 ± 0.6 for Rpn13-pru; Figure 4C; **Supporting Figure S4**). One potential explanation is that Rpn13 acts as part of a “latch” mechanism, keeping the proteasome in a low-unfolding ability state until ubiquitin binding during substrate engagement releases the latch, de-repressing the proteasome. This may in part explain why substrates that lack ubiquitin are unable to activate the proteasome’s unfolding ability.

### Keap1 ubiquitination leads to branched ubiquitin chains and DHFR mono- and di-ubiquitination

Our unfolding ability results suggested that Keap1-ubiquitinated substrates might interact with two receptors while Rsp5-ubiquitinated substrates only interacted with a single receptor to active the proteasome’s unfolding ability. We had previously shown that Rsp5 ubiquitination gave K63-linked chains, while Keap1 ubiquitination gave mixed linkage chains, but details of the chain architecture were unclear ^22^. To better understand how chain linkage might influence unfolding ability, we first characterized ubiquitinated substrates using tandem mass spectrometry. A construct containing a GST-tag followed by the Neh2Dual degron was ubiquitinated, bound to GST beads, washed to remove free ubiquitin and any unanchored chains that were produced (as well as any other ubiquitinated proteins), and then the degron was cleaved and eluted using HRV 3C protease (Figure 5A, B). The purified protein was then subjected to a middle-down mass spectrometry approach wherein limited tryptic digestion was used to cleave before the terminal Gly-Gly on ubiquitin generating ubiquitin(1-74) monomers with Gly-Gly adducts to the lysines where linkage occured (Figure 5C) ^50,51^. MS1 characterization of the resulting ubiquitin monomer products showed that while the Rsp5-ubiquitinated substrate produced only ubiquitin(1-74) monomers lacking a GG modification (the distal ubiquitin) and monomers containing a single GG modification (ubiquitin within a chain), the Keap1-ubiquitinated substrate produced a large extent of doubly- and even triply-modified ubiquitin(1-74) monomers, indicating a high degree of branching (Figure 5D). MS2 analysis using UVPD indicated that the Rsp5-ubiquitinated substrate was, as expected, most consistent with K63-linked chains, while the Keap1-ubiquitinated substrate was consistent with the presence of both K48- and K63-linked chains.

**Figure 5.**
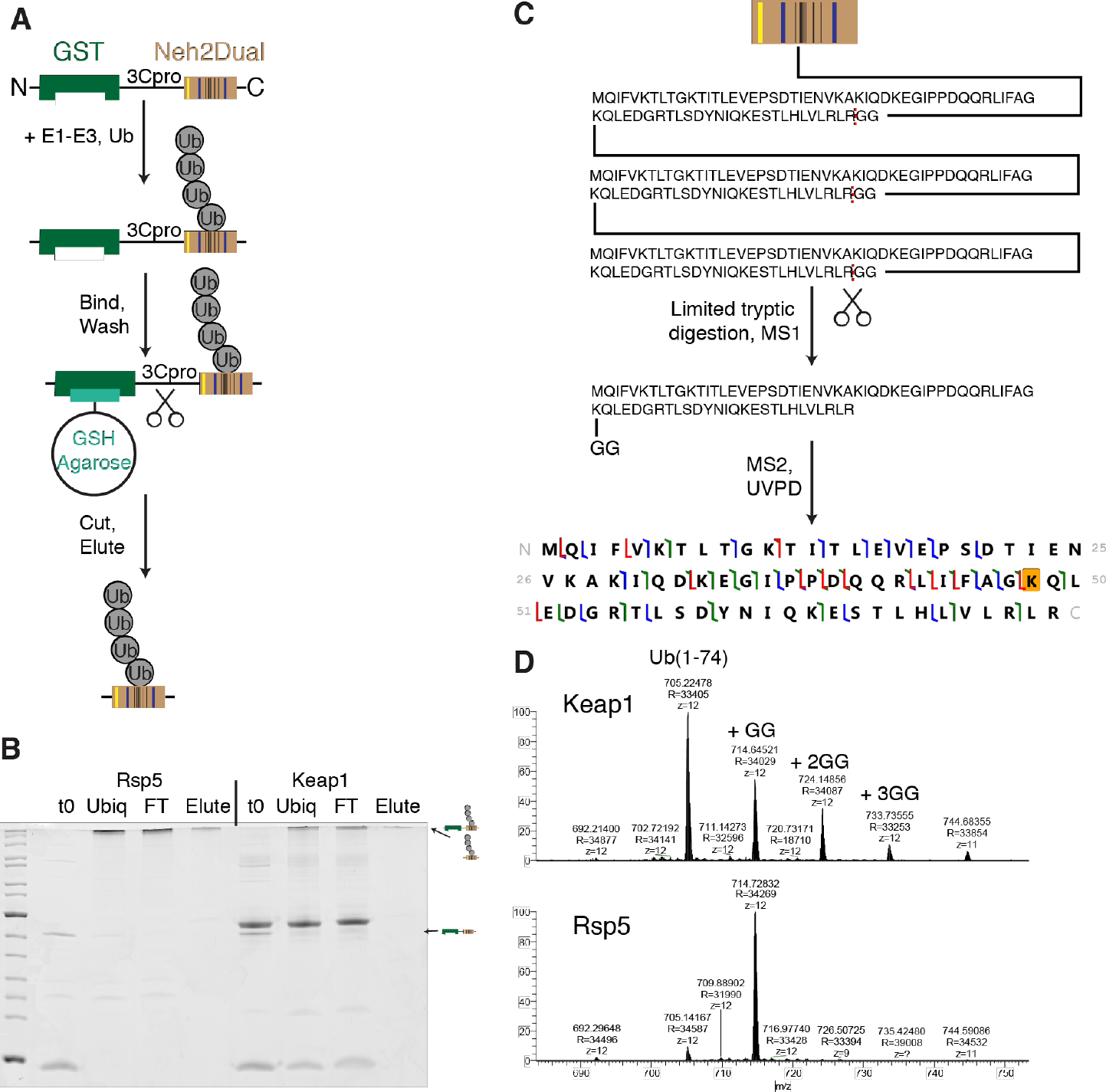
Characterization of Rsp5- and Keap1-ubiquitinated substrates. **A)**Schematic depiction of GST-Neh2dual substrate preparation. Yellow bar indicates Rsp5 binding site, blue bars Keap1 binding sites and black bars lysines in the Neh2 domain. Substrate was ubiquitinated with either Rsp5 or Keap1, bound to GSH-agarose beads, washed to remove free ubiquitin chains and components of the ubiquitination reaction, and then eluted by cleavage with GST-tagged HRV 3C protease. **B**) Coomassie-stained gel showing ubiquitination (t0 is a sample taken immediately after the reaction was initiated by addition of ubiquitin) and purification of the GST-Neh2Dual sample. Final elutions contain almost exclusively polyubiquitinated Neh2Dual with no contaminating free ubiquitin. Strong band just above the full-length substrate in Keap1 lanes is ovalbumin, a component of the ubiquitination buffer. **C**) Schematic depiction of middle-down mass spectrometry approach. Substrate (shown here as Ub_3_ attached to one lysine in Neh2Dual) was subjected to limited tryptic digestion to cleave ubiquitin prior to glycine 75 to generate ubiquitin(1-74) monomers. Here the proximal or middle ubiquitin is shown after digestion; this ubiquitin contains a diglycine appended to K48 that was derived from the distal ubiquitin in the chain. The distal ubiquitin would generate a ubiquitin(1-74) monomer lacking any diglycine modifications. Following cleavage, MS1 was used to identify the number of glycine modifications, and MS2 analysis with UVPD was used to determine the most likely site(s) of modification. **D**) Ub(1-74) monomers detected for Keap1 and Rsp5. For Keap1, monomers containing between zero and three diglycines were detected, indicating substantial branching, while for Rsp5, only monomers lacking or containing a single diglycine were detected, indicating linear chains.

Substrates containing branched ubiquitin chains have previously been shown to be degraded more efficiently by the proteasome than those containing linear chains ^11,52^. It therefore seemed possible that branching in the Keap1-modified substrate might be responsible for the increased unfolding ability observed with this substrate. Alternatively, differences in the locations of ubiquitin modifications on the substrate or extent of ubiquitination might affect the observed differences in unfolding ability. Control experiments using Cy5-labeled substrate and Cy3-labeled ubiquitin showed that each substrate is modified with about 20-30 ubiquitins in total (**Supporting Figure S5**); previous experiments suggested that small differences in the extent of substrate ubiquitination did not affect measured unfolding abilities ^22^.

We therefore ubiquitinated Neh2Dual-Barnase-DHFR with either Rsp5 or Keap1 and subjected it to bottom-up mass spectrometry to identify ubiquitination sites (**Figure S6**). Essentially all of the substrate is ubiquitinated on one or more lysines in the Neh2Dual degron, with very little modification outside this region **(Supporting Figure S6, Supporting Table S3**). The most common positions of modification detected for both Rsp5 and Keap1 were K43 and K49, the two most-N-terminal lysines within the degron (**Supporting Table S3**), and about 30% of the K43-ubiquitinated peptides detected were also ubiquitinated on K49, indicating that both substrates are capable of being ubiquitinated on more than one lysine simultaneously; previous experiments using K0 ubiquitin had shown that 5-7 lysines can be used in each substrate ^22^. However, there was some indication of occasional ubiquitination on the DHFR domain and, although mass spec is not quantitative, it appeared there was more ubiquitination on DHFR with Keap1 (10 modified peptides vs 133 unmodified detected at high confidence) than Rsp5 (6 modified vs 182 unmodified). Importantly, internal DHFR ubiquitination could potentially allow the DHFR-containing fragment to be re-targeted to the proteasome after it was released, thereby increasing the observed unfolding ability without changing the mechanics of substrate unfolding.

To quantitatively assess the extent of ubiquitination on DHFR, we used radiolabeled Neh2Dual-BarnaseL89G-C3Pro-DHFR-His, which contains an HRV 3C protease cleavage site between barnase and DHFR (Figure 6A). Cleavage of this substrate after ubiquitination produced a DHFR-containing fragment, which was then repurified via the C-terminal His-tag. If any ubiquitinated DHFR was produced, treatment with the non-specific deubiquitinase vOTU ^24^ should deubiquitinate it, leading to an increase in the amount of DHFR fragment. For Rsp5, no ubiquitinated fragment was observed, and quantitation of four replicate assays showed there was only 6 ± 3% total ubiquitination on the DHFR domain. However, for Keap1 several faint bands corresponding to mono- and di-ubiquitination were observed in the purified fragment, and vOTU treatment revealed that ~30% of DHFR domains were ubiquitinated, largely via the observed mono- and di-ubiquitination. The ubiquitination on this DHFR fragment was not sufficient to allow proteasomal degradation, even if a 43-amino acid unstructured tail was provided between the 3C-pro site and the DHFR domain to allow efficient initiation ^16^ (Figure 6B,C), suggesting mono and di-ubiquitination of the DHFR domain by Keap1 do not allow released DHFR fragment to be re-targeted to the proteasome.

**Figure 6.**
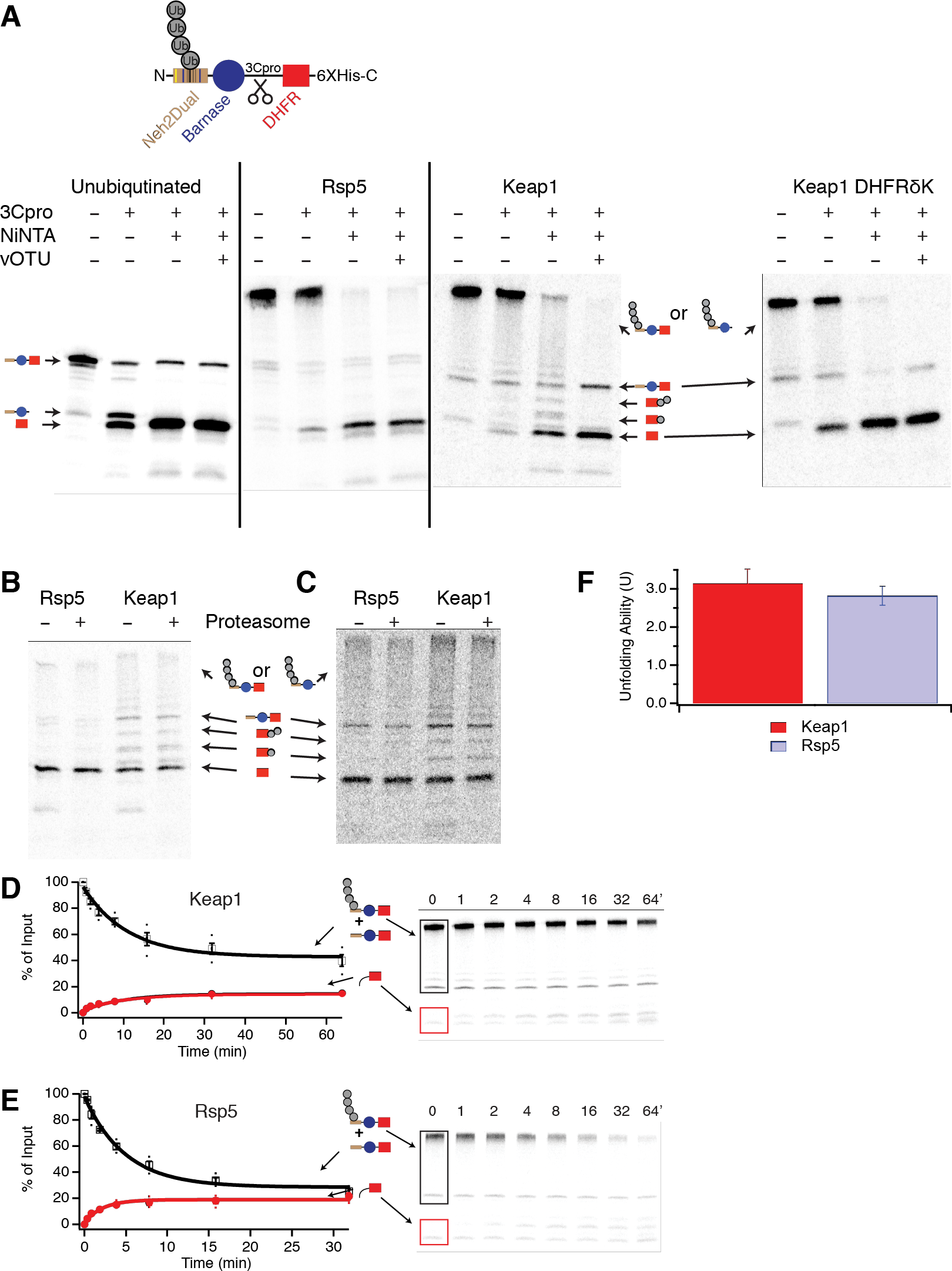
Substrate is polyubiquitinated on the degron, not the DHFR domain. **A**) Neh2Dual-BarnaseL89G-3Cpro-DHFR-His or Neh2Dual-BarnaseL89G-3Cpro-DHFRδK-His was ubiquitinated (or left unubiquitinated), cleaved with 3C-pro protease, and then repurified on NiNTA magnetic beads. After elution, the sample was either treated or mock-treated with vOTU to remove any ubiquitins on the DHFR domain. DHFRδK sample was ubiquitinated in the presence of 500 µM NADPH. Lines indicate separate gels. **B**) The purified DHFR fragment from A was incubated with 100 nM wild-type proteasome for 1 hr at 30 °C. There was no change in intensity of any of the DHFR bands, indicating that these DHFR fragments are not degraded or deubiquitinated by the proteasome. **C**) As in B, except substrate was Neh2Dual-BarnaseL89G-3Cpro-35ΔK-DHFR-6XHis, such that the purified fragment contained an unstructured initiation site. There was no change in intensity of any of the DHFR bands, indicating that these DHFR fragments are not degraded or deubiquitinated by the proteasome. Cleavage was less efficient with this construct, resulting in higher amounts of uncleaved material and less fragment. **D**, **E**) Degradation of trace radiolabeled Keap1-(**D**) or Rsp5- (**E**) ubiquitinated Neh2Dual-BarnaseL89G-3Cpro-DHFRδK-His by 100 nM wild-type proteasome. Black box shows region of the gel containing full-length protein with or without ubiquitination. Red box shows the region of the gel containing the DHFR fragment. The amounts of full-length protein (open squares) and DHFR fragment (red circles) are shown as a percentage of the ubiquitinated full-length substrate presented to the proteasome at the beginning of the reaction (or the full-length protein for the ubiquitin-independent substrate); the full-length protein is quantified as the sum of ubiquitinated and non-ubiquitinated full-length species so any deubiquitination is not misinterpreted as degradation. Dots are results from individual experiments, and error bars represent the SEM of 4 experiments. Curves are global fits to single exponentials. Sample gels are shown on the right. **F**) Unfolding abilities (U) calculated from the curve fits shown in **D** and **E**. Error bars are the SEM propagated from curve fitting the collected data sets.

The results of these mass spectroscopy and ubiquitination experiments suggest that Keap1-ubiquitinated substrates interact with multiple ubiquitin receptors either via branched chains on the degron or via poly-ubiquitination on the degron and mono or di-ubiquitination on the DHFR domain.

### Internal DHFR ubiquitination increases the proteasome’s unfolding ability but does not affect receptor utilization

To test whether branching or internal mono-ubiquitination was responsible for the higher unfolding ability seen with Keap1 than Rps5, we initially mutated all the lysines in DHFR to arginines. However, the resulting DHFR was so unstable that no DHFR containing fragment was ever released, even in the presence of NADPH or methotrexate (MTX) (data not shown). We therefore mutated individual lysines within DHFR in the Neh2Dual-Barnase-3Cpro-DHFR-His construct, and determined the extent of DHFR ubiquitination by purifying the DHFR-containing fragment after HRV 3C protease cleavage and treating with vOTU. Upon mutation of four lysines in DHFR (K32R/K58R/K106R/K109R; we refer to this DHFR mutant as DHFRδK) and ubiquitination in the presence of NADPH, no further mono- or di-ubiquitin bands were seen in the purified fragment and, based on quantification, no increase in fragment intensity was seen upon vOTU treatment (Figure 6A). The unfolding ability of each of these substrates, as measured by the extent of release of DHFR-containing fragment (Equation 1), was identical within error (Figure 6D-F), in contrast to previous experiments with wild-type DHFR in which the unfolding ability for Keap1 was approximately twice that of Rsp5 ^22 †^ (See also Figure 3). Thus, the internal mono- and di-ubiquitination on wild-type DHFR caused by Keap1 appears to increase the proteasome’s unfolding ability beyond ubiquitination on the degron alone, either by directly destabilizing DHFR or by allowing the substrate to continue to interact with ubiquitin receptors further into the degradation process, thereby keeping the proteasome in a fully active conformation longer. However, when we assayed the unfolding ability of the DHFRδK-containing substrate with ubiquitin receptor mutants (Figure 7, **Supporting Table S4**), we found an identical pattern of unfolding ability defects to the original lysine-containing substrate. For example, the Rpn1ΔT1/Rpn10ΔUIM double-mutant had a substantially reduced unfolding ability with the Keap1-ubiquitinated Neh2Dual-Barnase-3Cpro-DHFRδK-His substrate (but not with the Rsp5-ubiquitinated substrate), indicating that two ubiquitin receptors are still required for maximal unfolding ability with this substrate even without internal ubiquitination on the DHFR domain. Therefore, the architecture of polyubiquitination on the degron determines which receptors are required for activation of the proteasome’s unfolding ability.

**Figure 7.**
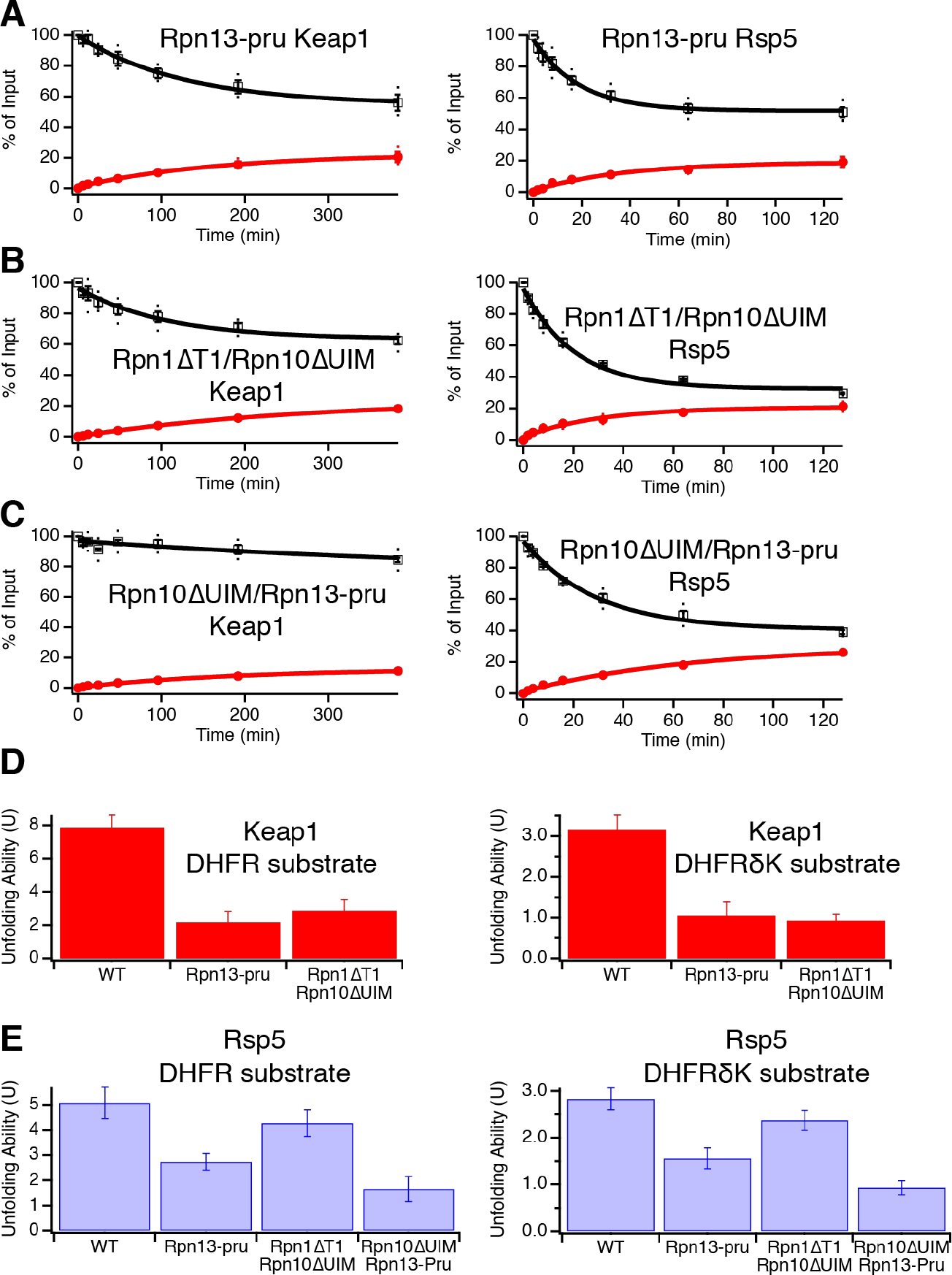
Ubiquitination on the DHFR domain does not affect differential dependence on ubiquitin receptors for Keap1 and Rsp5-ubiquitinated substrates. Degradation of trace radiolabeled Keap1- or Rsp5-ubiquitinated Neh2Dual-BarnaseL89G-3CPro-DHFRδK-His substrate by 100 nM Rpn13-pru (**A**), Rpn1ΔT1/Rpn10ΔUIM (**B**), or Rpn10ΔUIM/Rpn13-pru (**C**) proteasome. The amounts of full-length protein (open squares) and DHFR fragment (red circles) are shown as a percentage of the full-length substrate presented to the proteasome at the beginning of the reaction. Dots are results from individual experiments, and error bars represent the SEM of 3-4 experiments (except for Keap1 Rpn10ΔUIM/Rpn13-pru, which was 2 experiments and did not show appreciable degradation, so no U was calculated). Curves are global fits to single exponentials. **D, E**) Unfolding abilities (U) for Keap1 (**D**) and Rsp5 (**E**) substrates calculated from the curve fits shown in Figure 6 and A-C for the DHFRδK substrate compared to those for the DHFR containing substrate Neh2Dual-BarnaseL89G-DHFR-His from Figure 4. Error bars are the SEM propagated from curve fitting the collected data sets. The relative effects of each ubiquitin receptor mutation were very similar regardless of whether DHFR could be ubiquitinated.

## Discussion

It has long been known that the proteasome contains receptors that allow it to bind to ubiquitinated proteins. However, as the number of known receptors has increased, it has been puzzling why the proteasome requires at least three seemingly redundant receptors. In light of the recent discoveries that polyubiquitin modification not only targets proteins to the proteasome but also activates the proteasome’s peptidase activity ^53–55^, ATPase activity ^46^ and unfolding ability ^22^, it seemed that some subset of ubiquitin receptors might be involved in these activation processes. Our results indicate that Rpn13 is the major ubiquitin receptor responsible for recognizing ubiquitinated substrates, with both the largest effect on degradation rate and the largest effect on the proteasome’s unfolding ability upon mutating the ubiquitin binding site. Intriguingly, Rpn13 may serve to stabilize a low-unfolding ability conformation of the proteasome before ubiquitinated substrates bind (for example the S1 substrate-accepting state suggested by cryoEM ^56^). After a ubiquitinated substrate binds to Rpn13, a conformational change may occur alongside substrate engagement that stabilizes a high-unfolding ability state. Indeed, recent cryoEM structures solved in the presence of tetraubiquitin suggest that ubiquitin binding can cause changes in the conformation of the proteasome’s lid that are on the pathway towards activation and degradation ^57^. The continued presence of the substrate in the central channel of the 19S particle and its engagement with Rpt motor proteins may then maintain this activated state throughout the unfolding and degradation process, thereby potentially allowing the proteasome to “remember” that a polyubiquitin chain was attached to the substrate at the onset of degradation.

How might binding of polyubiquitin be communicated from the receptors to the Rpt motor proteins? Rpn13 is connected to the proteasome via attachment to a flexible extension at the C-terminus of Rpn2 (Figure 8A), and is fairly poorly resolved in cryoEM structures, suggesting that a range of conformations may be possible ^58^. Ubiquitin occupancy may then be communicated to the ring of Rpt motor proteins via these different conformations. In recent atomic-resolution cryoEM structures of the yeast proteasome ^59^, there is a different orientation of Rpn13 in the S1 substrate-accepting state versus the S3 putative translocating state, which seems to correlate with a substantial movement at the C-terminus of Rpn2 (~8 Å backbone movement at the last visible residue, Asp925, when Rpn2 is fixed and the structures are aligned) (Figure 8A). A nearby helical protrusion (787-821; shown in blue) has an even more dramatic change, with a rotation near the body of Rpn2 resulting in a ~15 Å distance change at the end of the helix. This helix in turn contacts the Rpt1/Rpt2 coiled coil, forming extensive contacts in the S1 state (potentially serving as a “latch” stabilizing this state) and more limited contacts in the S3 state, providing a potential pathway for communication between ubiquitin binding at Rpn13 and conformational change at the ATPases (Figure 8A). Indeed, in the archaeal homolog of the proteasome (PAN), conformational changes in the coiled coil structurally similar to Rpt1/Rpt2 have been implicated in controlling the activity of the PAN ATPases ^60^.

**Figure 8.**
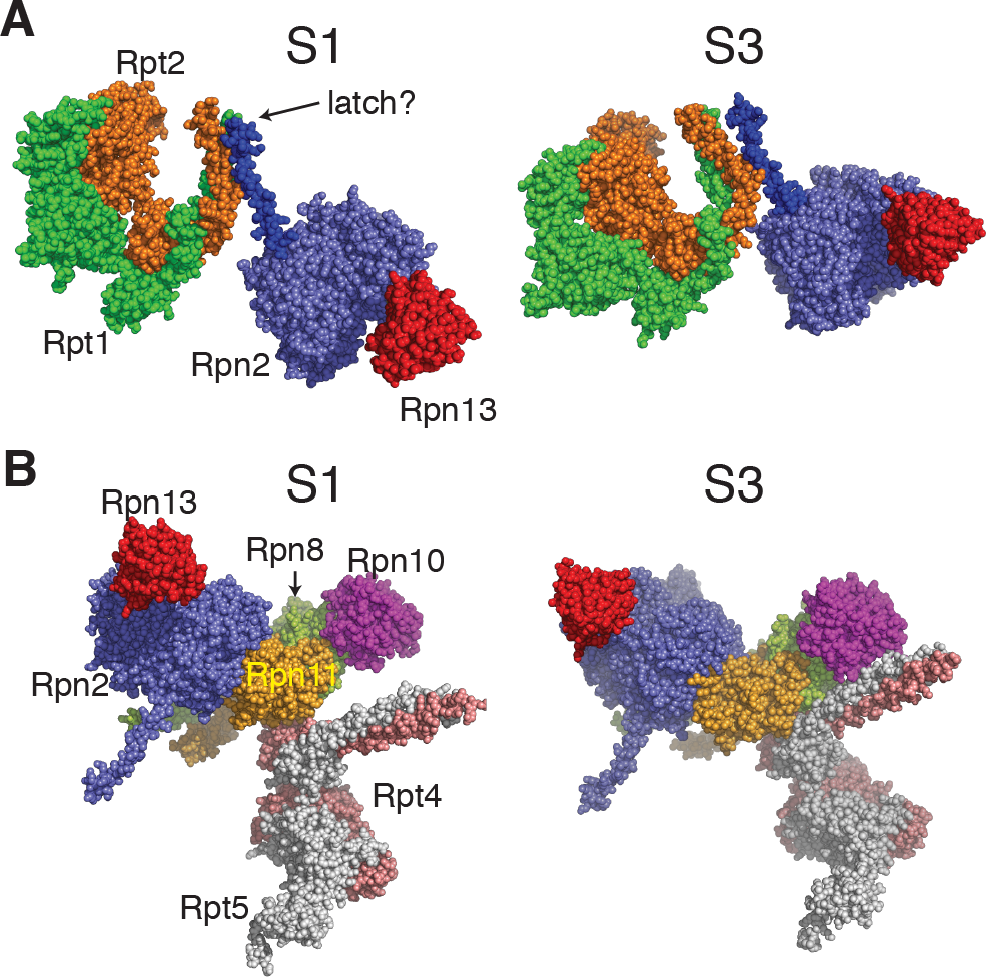
Conformational changes that could link ubiquitin binding to proteasomal activation. Left is S1 state (PDB IDs 5MP9 & 5MPD), right is S3 state (PDB ID 5MPB) **A)** Reorientation of Rpn13 (red) is correlated with movement of an Rpn2 (purple) helix (blue) to minimize contact with the Rpt1 (green)/Rpt2 (orange) coiled-coil. **B)** The Rpn8 (light green)/Rpn11 (light orange) heterodimer connects Rpn10 (magenta) and Rpn2. Rpn10 contacts the Rpt4 (salmon)/Rpt5 (grey) heterodimer in the S3 but not the S1 state.

Another pathway for communication might involve the Rpn8/Rpn11 heterodimer, which sits between Rpn2 and Rpn10, and has been shown to be in direct communication with the ATPase motors (Figure 8B) ^18^. Rpn10 interacts with the N-terminal coiled-coil of the Rpt4/Rpt5 pair in the S3 translocating state but not the S1 substrate-accepting state of the proteasome ^44^. Support for the involvement of Rpn10 in the pathway comes from the larger effect of removing the entire subunit on the proteasome’s unfolding ability than simply preventing ubiquitin binding.

Binding of ubiquitin to other receptors can also play a role in activating the proteasome’s unfolding ability. Our results suggest that the Rsp5-ubiquitinated substrate, perhaps due to its extended linear K63-linked chains, can interact with either Rpn13 or Rpn10 (but not Rpn1) to partially activate the proteasome, such that only mutation of both receptors dramatically reduces the unfolding ability. Presumably binding to either receptor can position the substrate such that the Rpt subunits can engage the substrate, but the geometry is unfavorable for simultaneous interaction with Rpn10, Rpn13 and the ATPases. Interestingly, our substrate was able to be degraded even in the absence of the Rpn10 UIM domain, while in previous work from the Martin lab, degradation of an Rsp5-ubiquitinated GFP-containing substrate was completely dependent on Rpn10 ^37^. Possibly this substrate cannot bind to Rpn13 in a productively positioned manner or has different unfolding requirements than our substrate, as the topology and stability of a substrate may affect the force needed to unfold it ^16,61,62^. Alternatively, reconstituted proteasome could be missing physiological modifications or proteasome-associated proteins (such as shuttle factors) that allow Rpn13 to be used productively or increase the proteasome’s unfolding ability.

For Keap1-ubiquitinated substrates, on the other hand, Rpn13 is necessary but insufficient for activation, suggesting that the branched ubiquitin modification allows simultaneous engagement of Rpn13, Rpn10 or Rpn1 and the ATPases and that internal mono- or di-ubiquitination on the DHFR may allow continued engagement of an ancillary receptor during the engagement and/or pulling process, giving a larger unfolding ability. Alternatively, the documented preferences of Rpn1, Rpn10 and Rpn13 for K48 versus K63-linked chains could play a role in allowing simultaneous engagement at multiple receptors ^14,63,64^.

More work is required to determine the mechanism by which binding of ubiquitinated substrates to Rpn13 and other ubiquitin receptors (including ubiquitin shuttle proteins) influences the unfolding ability of the proteasome, and the combinations of substrate and polyubiquitin geometries and architectures that maximize or minimize the proteasome’s ability to unfold its substrates.

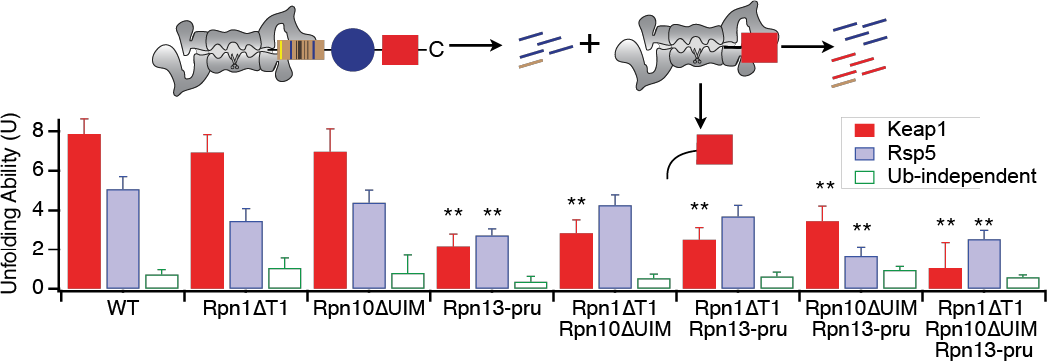

## Acknowledgements

The authors thank Aimee Eggler, Jennifer Palenchar, Barry Selinsky and Dennis Wykoff for helpful discussions. This material is based upon work supported by the National Science Foundation under Grant No. 1515229 to DAK. Support from NIH (R21CA191664 to JSB) and the Welch Foundation (F-1155 to JSB) is gratefully acknowledged. Funding from the UT System for support of the UT System Proteomics Core Facility Network is gratefully acknowledged.

## Abbreviations

5-FOA: 5-fluoroorotic acid
BSA: bovine serum albumin
DHFR: dihydrofolate reductase
DUB: deubiquitinase
pru: pleckstrin-like receptor for ubiquitin
MBP: maltose binding protein
MTX: methotrexate
UIM: ubiquitin interaction motif
UPS: Ubiquitin-Proteasome System
UVPD: ultraviolet photodissociation

## Supporting Information

List of yeast strains (**Table S1**), native gels of proteasome mutants (**Figure S1**), demonstration of directionality of degradation (**Figure S2**), degradation assays in the absence of NADPH (**Figure S3**), kinetic data and plots for additional assays (**Figure S4, Table S2,S4**) characterization of position and extent of ubiquitination of substrates (**Figure S5-S6, Table S3**).

## Footnotes

* The double-mutant (Rpn1ΔT1/Rpn13-pru) had an unfolding ability that was intermediate to but not significantly different from the Rpn13-pru single mutant (p = 0.14) or from wild-type (p = 0.10).

† The absolute unfolding abilities were also lower with the DHFRδK-containing substrate, but given that we have both changed the DHFR domain and the sequence the proteasome interacts with when unfolding the DHFR domain, this change is difficult to interpret.

