## Supporting Information for "Ubiquitin receptors are required for substrate-mediated activation of the proteasome’s unfolding ability"

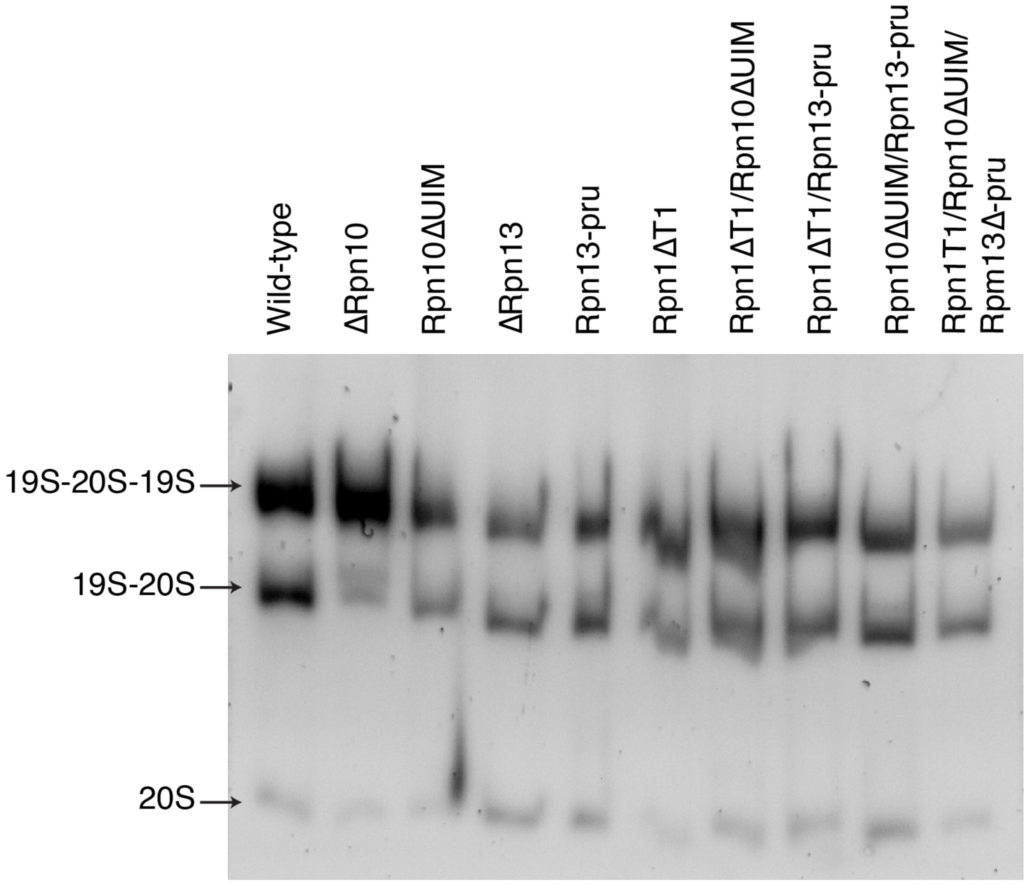


**Supporting Figure S1**. Native gel analysis of proteasome preps. ~6 μg of each proteasome prep was run on a 3.5% native gel, followed by visualization in the presence of 50 μM Suc-LLVY-AMC and 0.02% SDS.

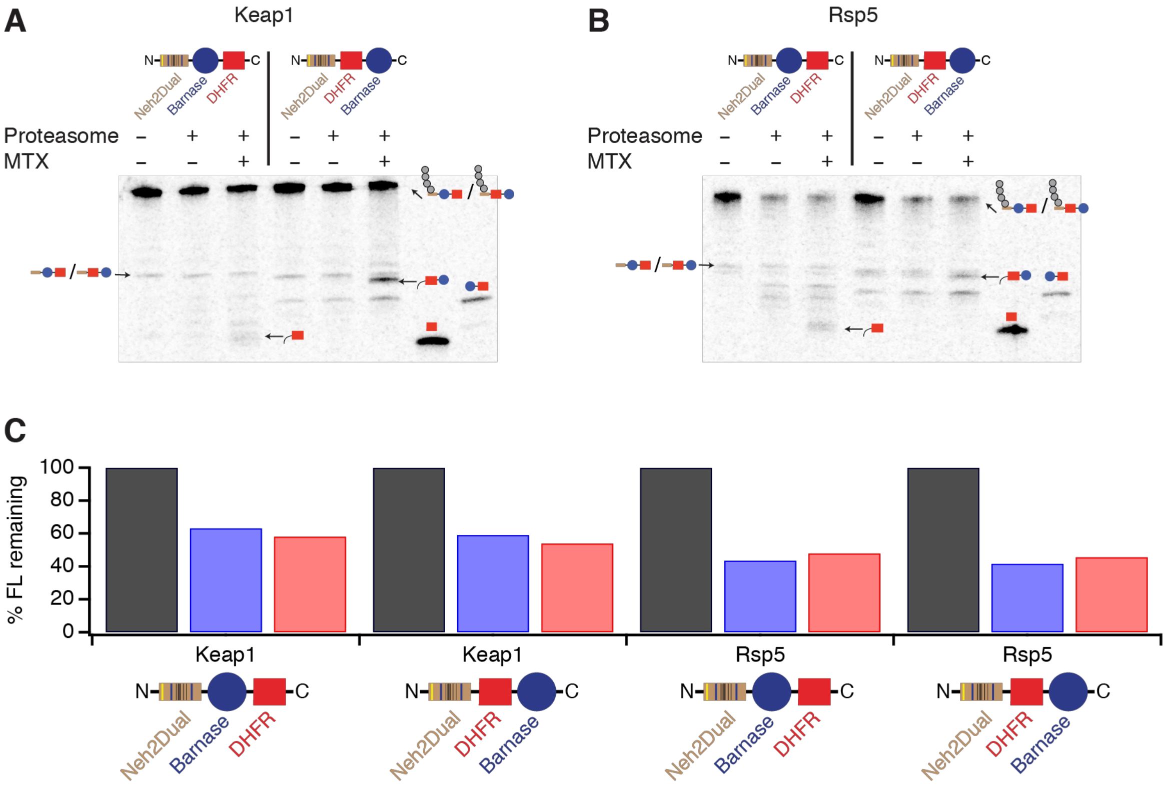


**Supporting Figure S2**. Degradation proceeds from the N-terminal degron towards the C-terminus. **A,B**) Degradation of trace labeled Keap1- (**A**) or Rsp5- (**B**) ubiquitinated Neh2Dual-Barnase-DHFR or Neh2Dual-DHFR-Barnase by 100 nM wild-type proteasome in the presence or absence of 100 µM MTX (2 hours). Fragments containing either DHFR plus a small tail or DHFR-Barnase plus a small tail were formed in the presence of proteasome and MTX from the Neh2Dual-Barnase-DHFR and Neh2Dual-DHFR-Barnase constructs respectively, and were identified relative to size standard control constructs (Barnase-DHFR and DHFR). **C**) Quantification of extent of degradation of ubiquitinated substrates in A and B. Grey bars are no proteasome, blue is proteasome only, red is proteasome plus MTX.

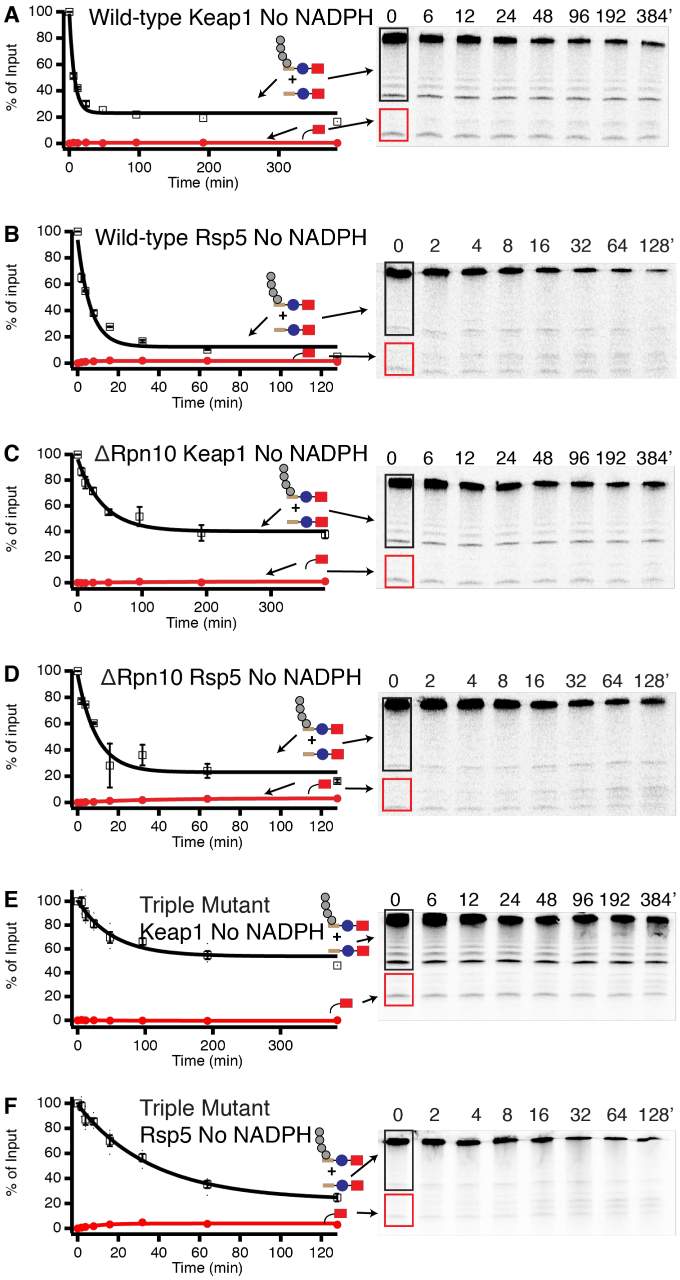


**Supporting Figure S3**. Proteasome mutants can degrade weakly folded proteins. Degradation of trace radiolabeled Keap1- or Rsp5-ubiquitinated Neh2Dual-BarnaseL89G-DHFR substrate by 100 nM wild-type (**A**,**B**), ΔRpn10 (**C**,**D**), or triple-mutant ubiquitin receptor (**E**,**F**) proteasome (Rpn1ΔT1/Rpn10ΔUIM/Rpn13-pru). With a substrate containing a destabilized barnase domain and in the absence of NADPH, degradation occurs without or with minimal fragment formation (amplitude for fragment formation with Rsp5-ubiquitinated substrate with wild-type is 1.7 ± 0.4%, with ΔRpn10 is 3.3 ± 0.3%, and with the triple-mutant is 4.0 ± 0.5%; no fragment was observed with Keap1-ubiquitinated substrates for any of the proteasomes). The amounts of full-length protein (open squares) and DHFR fragment (red circles) are shown as a percentage of the full-length substrate presented to the proteasome at the beginning of the reaction. Dots are results from individual experiments, and error bars represent the SEM of 4 experiments. Curves are global fits to single exponentials.

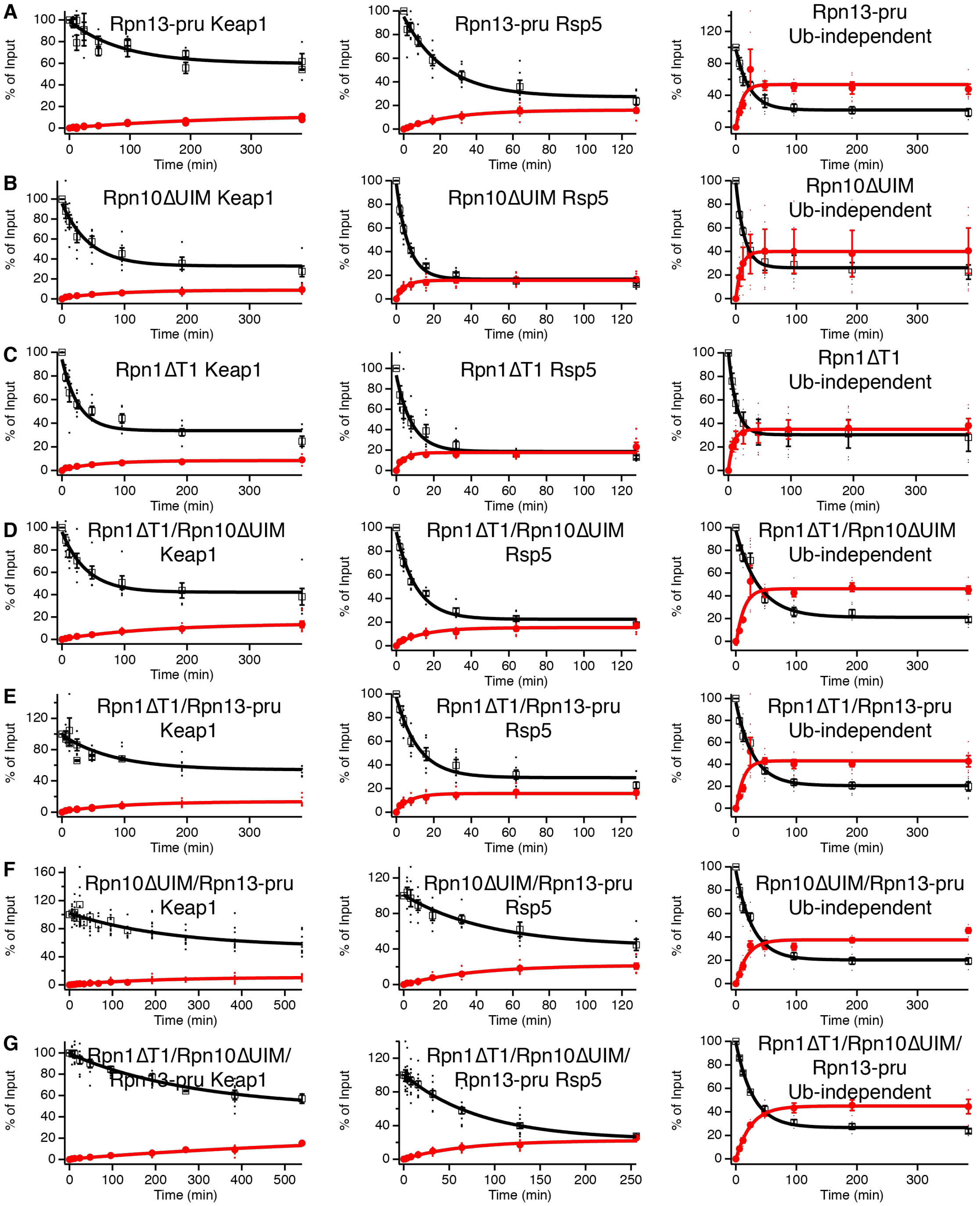


**Supporting Figure S4**. Degradation assays for mutant proteasomes. Degradation of trace radiolabeled Keap1-, Rsp5-ubiquitinated, or ubiquitin-independent substrate by 100 nM Rpn13-pru (**A**), Rpn10ΔUIM (**B**), Rpn1ΔT1 (**C**), Rpn1ΔT1/Rpn10ΔUIM (**D**), Rpn1ΔT1/Rpn13-pru (**E**), Rpn10ΔUIM/Rpn13-pru (**F**), or Rpn1ΔT1/Rpn10ΔUIM/Rpn13-pru (**G**) proteasome. The amounts of full-length protein (open squares) and DHFR fragment (red circles) are shown as a percentage of the full-length substrate presented to the proteasome at the beginning of the reaction. Dots are results from individual experiments, and error bars represent the SEM of 4-15 experiments. Curves are global fits to single exponentials.

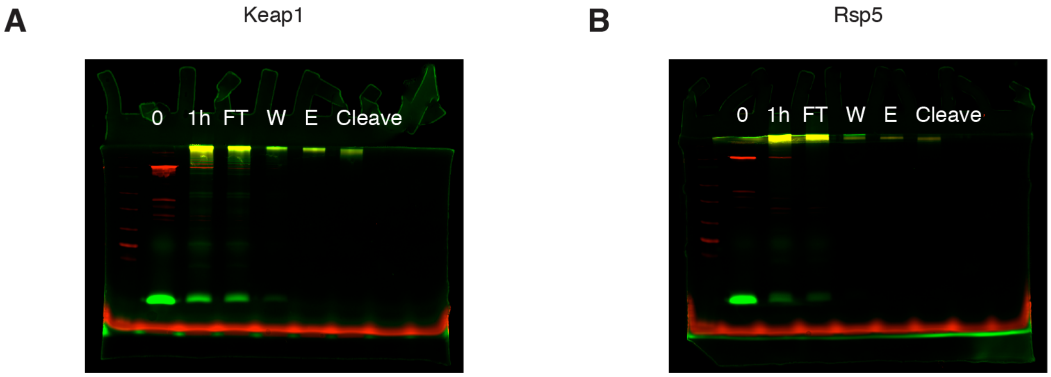


**Supporting Figure S5**. Estimation of number of ubiquitins per substrate. An MBP-3C$_{site}$-Barnase-Cys$^{Cy5}$-DHFR-His construct (red) was ubiquitinated with either **A**) Keap1 or **B**) Rsp5 for 1 hour in the presence of Cy3-labeled ubiquitin (green), bound to amylose resin, washed, eluted with maltose, and cleaved with HRV 3C protease to remove the MBP domain. The intensities of Cy3 and Cy5 in the final purified ubiquitinated substrate was compared to the intensities before ubiquitination or purification (and the known initial concentrations of substrate and ubiquitin) to determine the number of ubiquitins per substrate. Based on replicate experiments and comparisons to both the 0 and 1 hour timepoints, we estimate 30 ± 4 ubiquitins/Keap1 substrate and 22 ± 10 ubiquitins/Rsp5 substrate (the lower yield of the Rsp5 substrate makes estimation more difficult, but manual counting of individual ubiquitin bands at early time points using both fluorescent and radiolabeled substrates suggests that both Rsp5- and Keap1-ubiquitinated substrates carry in excess of 10 ubiquitins).

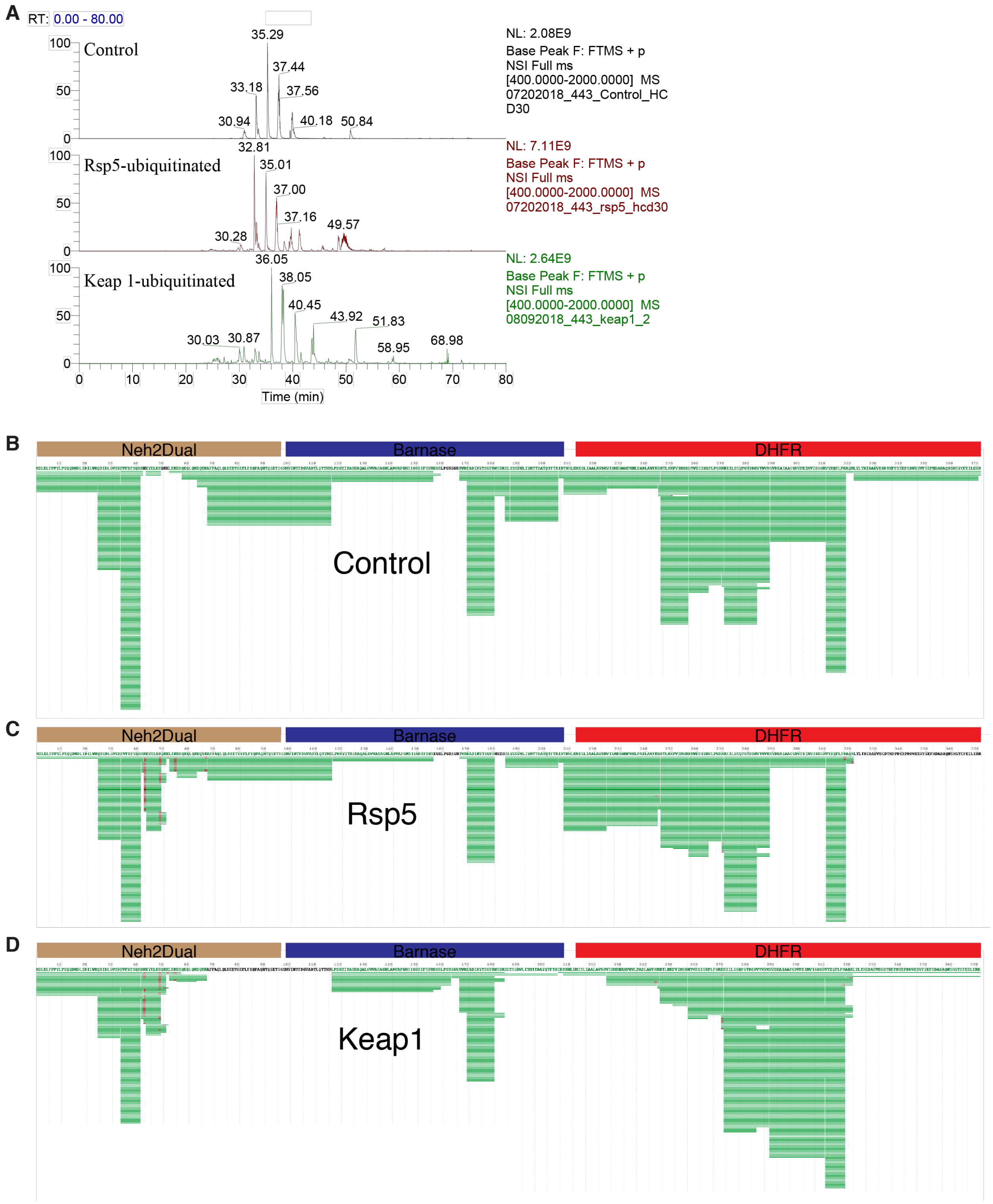


**Supporting Figure S6**. Bottom-up mass spectrometry of Neh2Dual-Barnase-DHFR identifies ubiquitination sites. **A**) Representative LC traces obtained for control (non-ubiquitinated) and ubiquitinated samples after tryptic digestion. **B-D**) Peptide coverage maps for **B**) control, **C**) Rsp5-ubiquitinated and **D**) Keap1-ubiquitinated samples. Green lines are peptides, red marks are sites of ubiquitination.

**Supporting Table S1**. Yeast strains. All strains derived from YYS40[27].

| Strain | Name | Genotype |
|---|---|---|
| YYS40 | Wild-type | MATa RPN11-3FLAG::HIS3 |
| yNDN1 | ΔRpn10 | MATa RPN11-3FLAG::HIS3ΔRpn10::NatMX |
| yNDN2 | ΔRpn13 | MATa RPN11-3FLAG::HIS3ΔRpn13::NatMX |
| yDAK36 | Rpn1ΔT1 | MATa RPN11-3FLAG::HIS3 ΔRpn1::NatMX Rpn1ΔT1::LEU2 (cen plasmid) |
| yDAK34 | Rpn10ΔUIM | MATa RPN11-3FLAG::HIS3 ΔRpn10::NatMX Rpn10ΔUIM::LEU2 (cen plasmid) |
| yMDC3 | Rpn13-pru | MATa RPN11-3FLAG::HIS3 ΔRpn13::NatMX Rpn13-pru::URA3 (cen plasmid) |
| yDAK39 | Rpn1ΔT1/Rpn10ΔUIM | MATa RPN11-3FLAG::HIS3 ΔRpn1::KanMX ΔRpn10::NatMX Rpn1ΔT1::LEU2 (cen plasmid) Rpn10ΔUIM::URA3 (cen plasmid) |
| yDAK44 | Rpn1ΔT1/Rpn13-pru | MATa RPN11-3FLAG::HIS3 ΔRpn1::KanMX ΔRpn13::NatMX Rpn1ΔT1::LEU2 (cen plasmid) Rpn13ΔUIM::URA3 (cen plasmid) |
| yDAK45 | Rpn10ΔUIM/ Rpn13-pru | MATa RPN11-3FLAG::HIS3 ΔRpn10::KanMX ΔRpn13::NatMX Rpn10ΔUIM::LEU2 (cen plasmid) Rpn13-pru::URA3 (cen plasmid) |
| yDAK47 | Rpn1ΔT1/Rpn10ΔUIM/ Rpn13-pru | MATa RPN11-3FLAG::HIS3 ΔRpn1::KanMX ΔRpn13::NatMX Rpn1ΔT1::LEU2 (cen plasmid) Rpn13ΔUIM::URA3 (cen plasmid) Rpn10ΔUIM |

**Supporting Table S2**. Summary of observed rate constants and unfolding abilities of wild-type and mutant proteasomes with Neh2Dual-BarnaseL89G-DHFR-His substrate. Errors are standard deviations propagated from global fitting.

| | Keap1-Ub | | Rsp5-Ub | | Ub-independent | |
|---|---|---|---|---|---|---|
| Mutant | $k_{obs}$ ($min^{-1}$) | U | $k_{obs}$ ($min^{-1}$) | U | $k_{obs}$ ($min^{-1}$) | U |
| WT | 0.051 ± 0.006 | 7.9 ± 0.8 | 0.14 ± 0.01 | 5.1 ± 0.6 | 0.033 ± 0.004 | 0.7 ± 0.2 |
| ΔRpn10 | 0.012 ± 0.007 | 4 ± 2 | 0.03 ± 0.01 | 2.5 ± 0.6 | 0.024 ± 0.006 | 0.5 ± 0.3 |
| ΔRpn13 | 0.020 ± 0.003 | 5.4 ± 0.6 | 0.05 ± 0.01 | 3.6 ± 0.5 | 0.027 ± 0.003 | 0.7 ± 0.3 |
| Rpn1ΔT1 | 0.038 ± 0.008 | 6.9 ± 0.9 | 0.12 ± 0.02 | 3.5 ± 0.6 | 0.08 ± 0.02 | 1.1 ± 0.5 |
| Rpn10ΔUIM | 0.023 ± 0.006 | 7 ± 1 | 0.16 ± 0.01 | 4.4 ± 0.6 | 0.07 ± 0.01 | 0.8 ± 0.9 |
| Rpn13-pru | 0.011 ± 0.004 | 2.2 ± 0.6 | 0.048 ± 0.007 | 2.7 ± 0.3 | 0.045 ± 0.008 | 0.4 ± 0.3 |
| Rpn1ΔT1/ Rpn10ΔUIM | 0.025 ± 0.007 | 2.9 ± 0.7 | 0.094 ± 0.009 | 4.3 ± 0.5 | 0.027 ± 0.004 | 0.5 ± 0.2 |
| Rpn1ΔT1/ Rpn13-pru | 0.012 ± 0.004 | 2.5 ± 0.6 | 0.08 ± 0.01 | 3.7 ± 0.6 | 0.035 ± 0.04 | 0.6 ± 0.2 |
| Rpn10ΔUIM/ Rpn13-pru | 0.004 ± 0.001 | 3.5 ± 0.8 | 0.02 ± 0.01 | 1.7 ± 0.5 | 0.036 ± 0.004 | 1.0 ± 0.2 |
| Rpn10ΔUIM/ Rpn13-pru* | 0.007 ± 0.004 | 4 ± 1 | 0.05 ± 0.01 | 2.2 ± 0.2 | ND | ND |
| Rpn1ΔT1/ Rpn10ΔUIM/ Rpn13-pru | 0.003 ± 0.001 | 1 ± 1 | 0.012 ± 0.003 | 2.6 ± 0.4 | 0.035 ± 0.002 | 0.6 ± 0.1 |

*Additionally purified via superose 6 gel filtration; data from 2 trials.

**Supporting Table S3**. Number of modified and unmodified peptides containing lysine residues identified for Rsp5- and Keap1-ubiquitinated Neh2Dual-Barnase-DHFR substrate. (Only peptides with PEP2D score lower than 0.001 were considered).

| **Rsp5** | **No. of modified peptides** | **No. of unmodified peptides** | **Domain** |
|---|---|---|---|
| K43 | 32 | 2 | Neh2Dual |
| K49 | 19 | 27 | Neh2Dual |
| K52 | 1 | 0 | Neh2Dual |
| K55 | 5 | 0 | Neh2Dual |
| K67 | 4 | 5 | Neh2Dual |
| K244 | 1 | 52 | DHFR |
| K270 | 2 | 11 | DHFR |
| K318 | 3 | 119 | DHFR |
| **Keap1** | **No. of modified peptides** | **No. of unmodified peptides** | **Domain** |
| K43 | 24 | 9 | Neh2Dual |
| K49 | 13 | 28 | Neh2Dual |
| K244 | 3 | 11 | DHFR |
| K270 | 5 | 27 | DHFR |
| K318 | 2 | 95 | DHFR |

**Supporting Table S4**. Summary of observed rate constants and unfolding abilities of wild-type and mutant proteasomes with Neh2Dual-BarnaseL89G-3CPro-DHFRδK-His substrate. Errors are standard deviations propagated from global fits.

| | Keap1-Ub | | Rsp5-Ub | |
|---|---|---|---|---|
| Mutant | $k_{obs}$ ($min^{-1}$) | U | $k_{obs}$ ($min^{-1}$) | U |
| WT | 0.09 ± 0.02 | 3.2 ± 0.4 | 0.21 ± 0.02 | 2.8 ± 0.2 |
| Rpn13-pru | 0.008 ± 0.002 | 1.1 ± 0.3 | 0.026 ± 0.004 | 1.6 ± 0.2 |
| Rpn1ΔT1/ Rpn10ΔUIM | 0.010 ± 0.004 | 0.4 ± 0.2 | 0.049 ± 0.004 | 2.4 ± 0.2 |
| Rpn10ΔUIM/ Rpn13-pru | < 0.01 | ND | 0.035 ± 0.006 | 0.9 ± 0.2 |
